# Correct CYFIP1 dosage is essential for synaptic inhibition and the excitatory / inhibitory balance

**DOI:** 10.1101/303446

**Authors:** Elizabeth C. Davenport, Blanka Szulc, James Drew, James Taylor, Toby Morgan, Nathalie F. Higgs, Guillermo Lopez-Domenech, Josef T. Kittler

## Abstract

Altered excitatory/inhibitory balance is implicated in neuropsychiatric disorders but the genetic aetiology of this is still poorly understood. Copy number variations in CYFIP1 are associated with autism, schizophrenia and intellectual disability but the role of CYFIP1 in regulating synaptic inhibition or excitatory/inhibitory balance remains unclear. We show, CYFIP1, and its paralogue CYFIP2, are enriched at inhibitory postsynaptic sites. While upregulation of CYFIP1 or CYFIP2 increased excitatory synapse number and the frequency of miniature excitatory postsynaptic currents (mEPSCs), it had the opposite effect at inhibitory synapses, decreasing their size and the amplitude of miniature inhibitory postsynaptic currents (mIPSCs). Contrary to CYFIP1 upregulation, its loss *in vivo*, upon conditional knockout in neocortical principal cells, increased expression of postsynaptic GABA_A_ receptor β2/3-subunits and neuroligin 3 and enhanced synaptic inhibition. Thus, CYFIP1 dosage can bi-directionally impact inhibitory synaptic structure and function, potentially leading to altered excitatory/inhibitory balance and circuit dysfunction in CYFIP1-associated neurodevelopmental disorders.

## Introduction

Schizophrenia (SCZ) and autism spectrum disorder (ASD) have a strong genetic component with a growing number of rare variant mutations and copy number variations (CNVs; deletions and duplications) in functionally overlapping synaptic and neurodevelopmental gene sets linked to increased disease susceptibility (Bourgeron, 2015; Fromer et al., 2014; Iossifov et al., 2014; Marshall et al., 2017; De Rubeis et al., 2014). Identifying how neuronal connectivity is altered by these genetic lesions is crucial for understanding nervous system function and pathology. CNVs of the 15q11.2 region of the human genome are implicated in the development of neurological and neuropsychiatric conditions. 15q11.2 CNV loss is associated with SCZ (Marshall et al., 2017; Rees et al., 2014; Stefansson et al., 2008), while numerous reports have identified 15q11.2 duplications and deletions in individuals with ASD (Doornbos et al., 2009; Picinelli et al., 2016; Pinto et al., 2014; van der Zwaag et al., 2010) epilepsy and intellectual disability (de Kovel et al., 2010; Nebel et al., 2016; Vanlerberghe et al., 2015). 15q11.2 contains 4 genes (NIPA1, NIPA2, CYFIP1 and TUBGCP5) with substantial evidence from rodent and human models pointing towards CYFIP1 as the main disease-causing gene within the locus (Bozdagi et al., 2012; Nebel et al., 2016; Oguro-Ando et al., 2014; Pathania et al., 2014; De Rubeis et al., 2013; Yoon et al., 2014). Polymorphisms and rare variants in CYFIP1 are also linked to susceptibility in ASD (Toma et al., 2014; Wang et al., 2015) and SCZ (Tam et al., 2010; Yoon et al., 2014) with a direct deletion of *CYFIP1* identified in an autistic patient with a *SHANK2* deletion (Leblond et al., 2012). Moreover, genome wide expression profiling of patients with a 15q11-13 duplication has demonstrated an up-regulation of *CYFIP1* mRNA in those that suffer from ASD, highlighting the importance of investigating the effects of genetic duplication as well as deletion (Nishimura et al., 2007). The CYFIP1 paralogue, CYFIP2, has also been linked to neurological disorders including SCZ, epilepsy, eating disorders, Alzheimer’s disease, Fragile X syndrome-like behaviours and cocaine seeking (Föcking et al., 2014; Han et al., 2015; Kirkpatrick et al., 2017; Kumar et al., 2013; Nakashima et al., 2018; Tiwari et al., 2016).

CYFIP1 and CYFIP2, are key components of the <u>W</u>AVE <u>R</u>egulatory <u>C</u>omplex (WRC; a hetero-pentamer consisting of WAVE, Abi, Nap1, HSPC300 and CYFIP1 or CYFIP2) that plays a critical role in regulating the dynamics of the actin cytoskeleton in cells by activating ARP2/3-mediated F-actin branching (Chen et al., 2010). Rare variants of Nap1 (NCKAP1) are also genetically linked to ASD and intellectual disability (Anazi et al., 2017; Iossifov et al., 2014; De Rubeis et al., 2014) providing further genetic support for a critical role of WRC-dependent actin regulatory pathways in neurodevelopmental disorders. Additionally, CYFIP1 is also a repressor of cap-dependent translation by acting as a non-canonical eIF4E binding protein in its complex with the ASD associated FMRP protein (Napoli et al., 2008) and can also modulate the mTOR pathway (Oguro-Ando et al., 2014).

Synaptic inhibition, mediated by GABA_A_ receptors (GABA_A_Rs), is vital for the efficient control of network excitability, excitation/inhibition (E/I) balance and for normal brain function. Inhibitory synapses require the stabilisation of postsynaptic GABA_A_Rs opposed to GABA-releasing presynaptic terminals. Modulation of inhibitory synaptic strength can be achieved by regulating the size and number of inhibitory synapses (Bannai et al., 2009; Luscher et al., 2011; Muir et al., 2010; Twelvetrees et al., 2010) and the clustering of GABA_A_Rs by an inhibitory postsynaptic complex containing the heteromeric scaffold gephyrin (Tyagarajan and Fritschy, 2014) and adhesion molecules such as neuroligins (Davenport et al., 2017; Pettem et al., 2013; Poulopoulos et al., 2009; Smith et al., 2014; Uezu et al., 2016; Yamasaki et al., 2017). While CYFIP1 is enriched at excitatory synapses where it can regulate F-actin dynamics (Pathania et al., 2014) and the development and plasticity of dendritic spines (Abekhoukh et al., 2017; Pathania et al., 2014; De Rubeis et al., 2013), the role of CYFIP1 at inhibitory synapses and in regulating the E/I balance remains undetermined.

Here we show that CYFIP1 and CYFIP2 are enriched at inhibitory synapses. CYFIP1 overexpression in dissociated neurons alters the excitatory to inhibitory synapse ratio, resulting in reduced mIPSC amplitude and increased mEPSC frequency. Conversely, when CYFIP1 is conditionally knocked-out from excitatory neocortical pyramidal cells, inhibitory synaptic components are upregulated and mIPSC amplitude is significantly increased. Thus, altered gene dosage of CYFIP1 disrupts inhibitory synaptic structure, leading to altered neuronal inhibition. Our data supports a role for CYFIP1 in regulating synapse number and the E/I balance and highlights a mechanism that may contribute to the neurological deficits observed in 15q11.2 CNV-associated neuropsychiatric conditions.

## Results

### CYFIP proteins are enriched at inhibitory synapses

While we and others have previously demonstrated an enrichment of CYFIP proteins at excitatory synapses (Pathania et al., 2014; De Rubeis et al., 2013), nothing is known regarding their localisation to inhibitory synapses. We therefore used immunofluorescence and confocal imaging to examine CYFIP1 and CYFIP2 subcellular distribution in cultured neurons. We found CYFIP1^GFP^ and CYFIP2^GFP^ exhibited a non-uniform distribution along dendrites appearing to be selectively targeted to punctate clusters in dendritic shafts in addition to the previously reported localisation of CYFIP1/2 to spine heads (Pathania et al., 2014) (**Fig. 1A,B**). Labelling neurons with antibodies to the inhibitory pre and postsynaptic markers VGAT and gephyrin respectively revealed that clusters of CYFIP1^GFP^ and CYFIP2^GFP^ in dendritic shafts could be found colocalised with gephyrin opposed to VGAT labelled inhibitory terminals (**Fig. 1A,B**). This can also be seen in the line scan of the zoom images where fluorescence intensity is plotted against distance. Indeed, quantitative image analysis revealed a ~40 % enrichment of CYFIP1^GFP^ and CYFIP2^GFP^ fluorescence at gephyrin clusters compared to the total process. We confirmed this result by labelling for endogenous CYFIP1 which was also found to be highly enriched at inhibitory synapses and colocalised with gephyrin clusters (**Fig. 1C**). To explore the distribution of CYFIP1 within inhibitory postsynaptic sites we carried out stimulated emission depletion (STED) microscopy to resolve this sub-synaptic compartment (Vicidomini et al., 2011). Interestingly, STED imaging performed on neurons labelled with antibodies to endogenous CYFIP1 and gephyrin revealed the presence of small CYFIP1 nanoclusters forming around gephyrin sub-synaptic domains (**Fig. 1D**). To further investigate the intimate association of CYFIP1 with the inhibitory postsynaptic scaffold we carried out a proximity ligation assay (PLA) (Norkett et al., 2015) on neurons labelled with antibodies to endogenous CYFIP1 and gephyrin. PLA detects interactions between endogenous proteins in fixed samples, giving a fluorescent readout after incubation with relevant primary antibodies, ligation, and amplification steps. The significant 2.7 fold increase in PLA puncta in the dual antibody condition indicated an intramolecular distance of <40 nm and demonstrates that CYFIP1 can complex with gephyrin in hippocampal neurons further supporting an enrichment of CYFIP1 at inhibitory postsynapses (**Fig. 1E,F**). These data indicate that CYFIP proteins can be found enriched at inhibitory synapses where they can intimately associate with the gephyrin scaffold.

**Figure 1:**
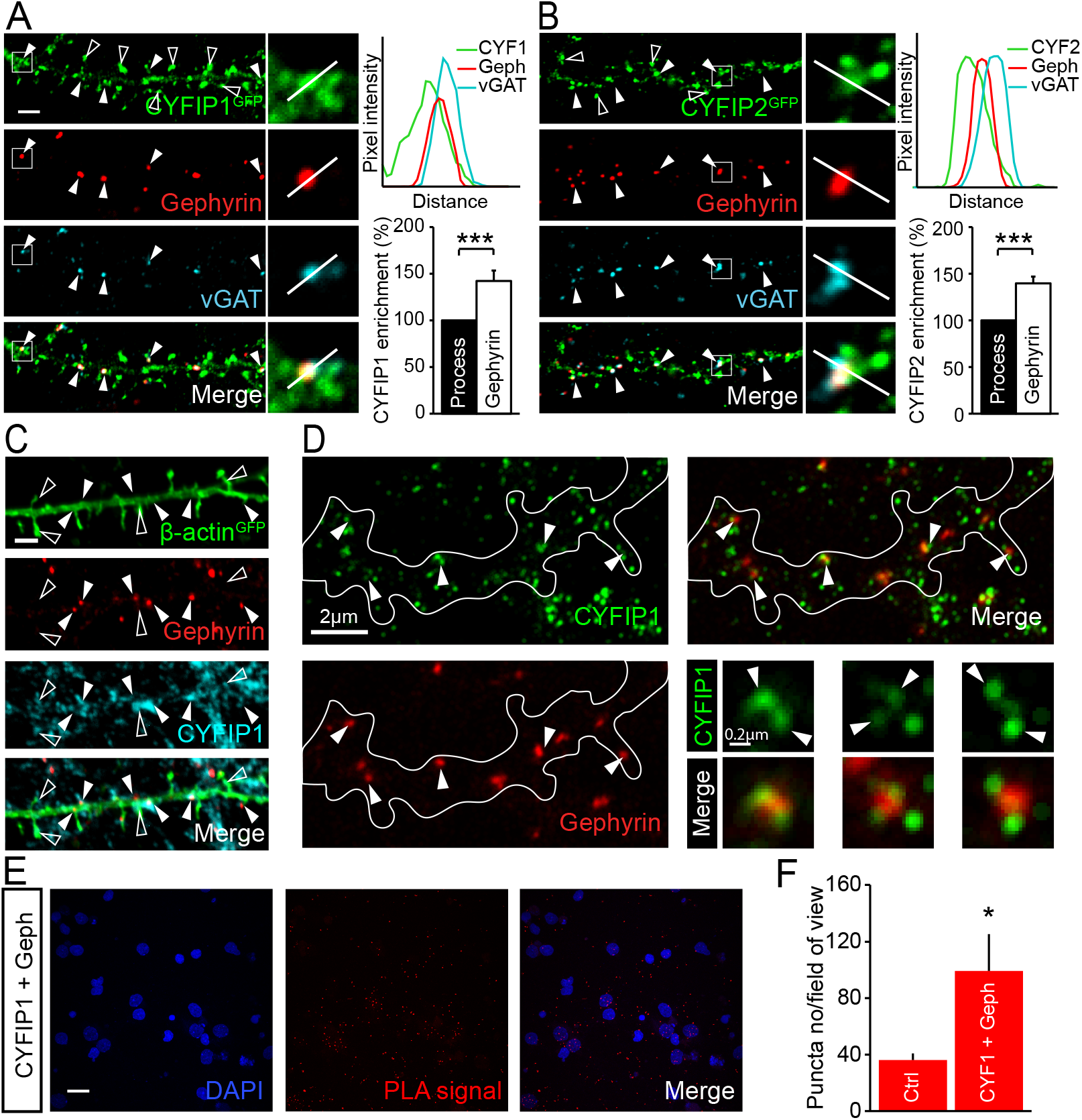
CYFIP1 and CYFIP2 are present at inhibitory synapses. **(A-B)** Confocal images show cultured hippocampal neurons transfected with CYFIP1^GFP^ (A) or CYFIP2^GFP^ **(B)** and immunolabelled for the inhibitory pre and postsynaptic markers VGAT and gephyrin respectively. CYFIP1^GFP^ and CYFIP2^GFP^ clusters colocalised with the inhibitory synaptic markers (arrowheads) and are also present at dendritic spines (open arrowheads). Graphs show line scans through clusters (top) and quantification of CYFIP1^GFP^ and CYFIP2^GFP^ fluorescence intensity at inhibitory synaptic gephyrin puncta compared to the total process (bottom) (CYFIP1: 42.4 ± 11.2% increase, p < 0.0001; CYFIP2: 39.8 ± 7.3% increase p = 0.0002; n = 33-42 processes from 9 cells from 3 independent preparations; Wilcoxon Signed Rank Test). Scale bar, 2 μm. **(C)** Endogenous CYFIP1 colocalises with the inhibitory postsynaptic marker gephyrin (filled arrowheads) and is also present in dendritic spines (open arrowheads) in hippocampal neurons transfected with the cell fill actin^GFP^. Scale bar, 2 μm. **(D)** STED images of endogenous CYFIP1 and gephyrin. Arrowheads show CYFIP1 nanoclusters at gephyrin puncta. Scale bar, 2 μm, zoom scale bar, 0.2 μm. **(E-F)** Example images and puncta quantification of proximity ligation assay (PLA) on hippocampal neurons using antibodies to CYFIP1 and gephyrin compared to single CYFIP1 antibody control conditions (control: 36.1 ± 4.7, CYFIP1 and gephyrin: 99.3 ± 26.1, n = 14 cells from 3 preparations, p = 0.0217; Mann-Whitney). Scale bar, 20 μm. * p < 0.05, ***p < 0.001. Bars indicate mean and error bars s.e.m.

### Upregulating CYFIP1 or CYFIP2 expression disrupts inhibitory synaptic structure and alters the excitatory to inhibitory synaptic ratio

Increased CYFIP1 copy number has been linked to neurodevelopmental alterations including ASD but the impact of increased CYFIP1 or CYFIP2 expression on synaptic function remains poorly understood. Given that we now show both proteins to be enriched at inhibitory synapses we investigated the impact of upregulating CYFIP1/2 expression on inhibitory synapse number and area. Cultured neurons were transfected for 4 days with CYFIP1^GFP^ or CYFIP2^GFP^ before being fixed at DIV14 and labelled with an antibody against gephyrin as a marker for inhibitory synapses. Quantification revealed that gephyrin cluster number and immunolabelled area was significantly reduced in both CYFIP1 and CYFIP2 overexpressing cells (**Fig. 2A,B**). Consistent with this, the total number and area of surface GABA_A_R clusters, labelled with an antibody raised to an extracellular epitope in the synaptically enriched GABA_A_R-γ2 subunit, were also significantly reduced (**Fig. 2C,D**).

**Figure 2:**
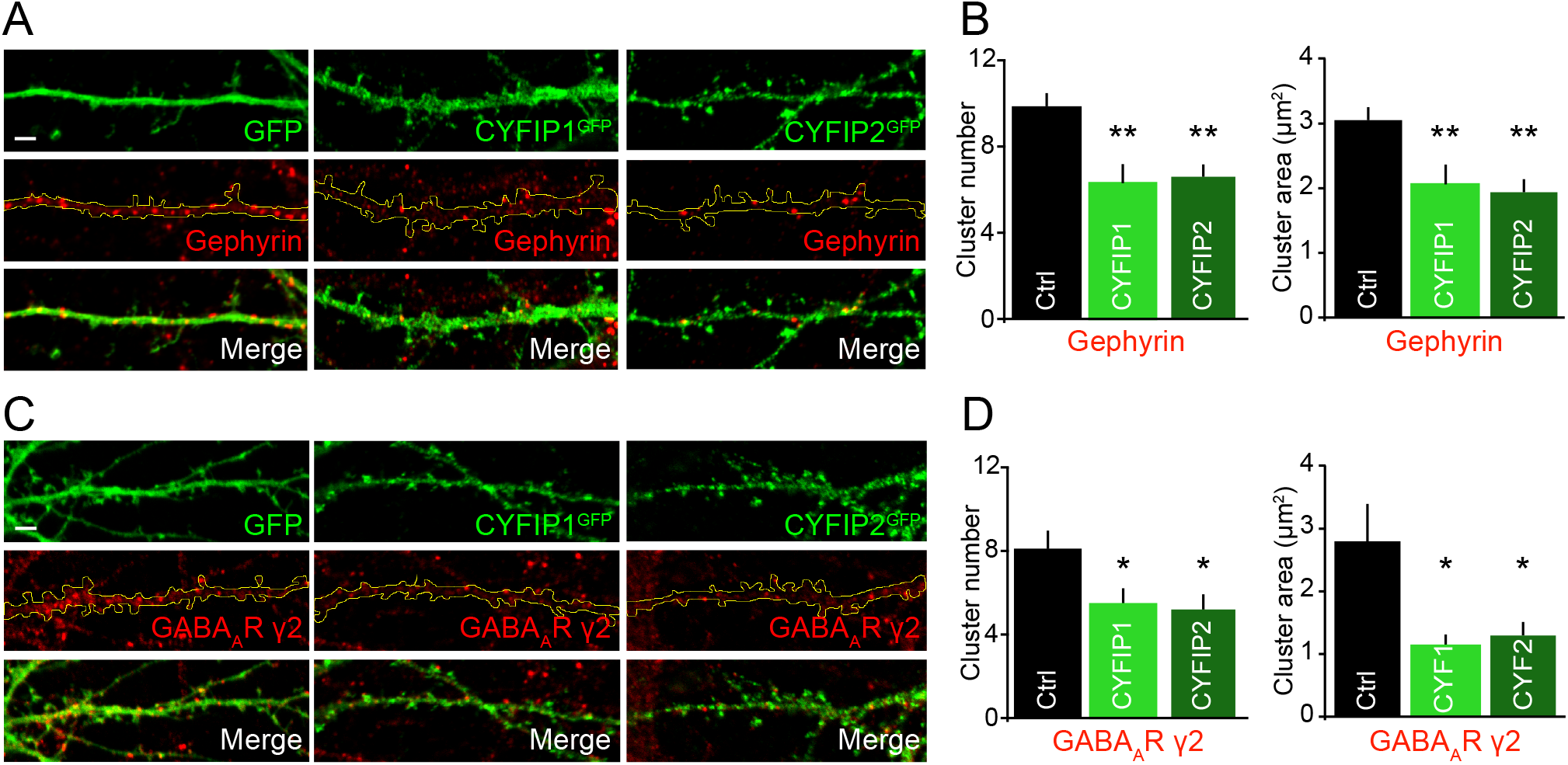
The effect of increased CYFIP1 and CYFIP2 gene dosage on inhibitory synaptic structure. **(A)** Representative confocal images of hippocampal neurons transfected with CYFIP1^GFP^, CYFIP2^GFP^ or GFP control for 4 days before fixing at DIV14 and labelling with an antibody to gephyrin. Scale bar, 2 μm. **(B)** Gephyrin cluster analysis showing a significant decrease in gephyrin cluster number and area upon CYFIP1^GFP^ or CYFIP2^GFP^ overexpression (cluster number: from 9.9 ± 0.6 to 6.3 ± 0.9 for CYFIP1 and 6.6 ± 0.6 for CYFIP2; cluster area: from 3.1 ± 0.2 μm^2^ to 2.1 ± 0.3 μm^2^ for CYFIP1 and 1.9 ± 0.2 μm^2^ for CYFIP2; n = 20 cells from 4 preparations; Kruskal-Wallis oneway ANOVA, Dunn’s post-hoc multiple comparisons). **(C)** Representative confocal images of hippocampal neurons transfected as (A) and surface labelled with an antibody to the GABA_A_R-γ2 subunit. Scale bar, 2 μm. **(D)** Cluster analysis of GABA_A_R-γ2 surface puncta showing a decrease in cluster number and area upon CYFIP1^GFP^ or CYFIP2^GFP^ overexpression (cluster number: from 8.1 ± 0.9 to 5.4 ± 0.8 for CYFIP1 and 5.2 ± 0.7 for CYFIP2; cluster area: from 2.8 ± 0.6 μm^2^ to 1.1 ± 0.2 μm^2^ for CYFIP1 and 1.2 ± 0.2 μm^2^ for CYFIP2; n = 25 cells from 4 preparations; Kruskal-Wallis oneway ANOVA, Dunn’s post-hoc multiple comparisons). *p < 0.05, **p < 0.01. Bars indicate mean and error bars s.e.m.

Remarkably, when neurons were labelled with an antibody to the excitatory PSD scaffold protein homer to label excitatory synapses the opposite effect was observed. Notably, the total number and area of homer clusters along dendrites was significantly increased in CYFIP1/2 overexpressing cells (**Fig. 3A,B**). To determine if the CYFIP1 overexpression-dependent increase in excitatory postsynapse number correlated with an increase in functional synapses we analysed the number of innervated excitatory synapses along the dendritic region, considered as the number of overlapping VGLUT-labelled presynaptic and PSD95-labelled postsynaptic puncta. Innervated synapses were significantly increased in cells overexpressing CYFIP1 compared to control and consistent with this the number of presynaptic VGLUT clusters were also enhanced (**Fig. 3C-E**). This alteration in excitatory synaptic number led us to investigate where within the dendrite these new synapses were forming. There were significantly more excitatory synapses on both the dendritic shaft and spines in CYFIP1 overexpressing cells compared to control which resulted in an increased proportion of the total number of synapses present on the shaft (**Fig. S2A-C**). Consistently, the ratio of synapses on the spine verses the shaft deceased in CYFIP1 overexpressing cells (**Fig. S2D**). Interestingly, there was no change in spine density along dendrites although spine morphology was altered with significantly more long thin and mushroom spines in cells overexpressing CYFIP proteins (**Fig. S2E-G**).

**Figure 3:**
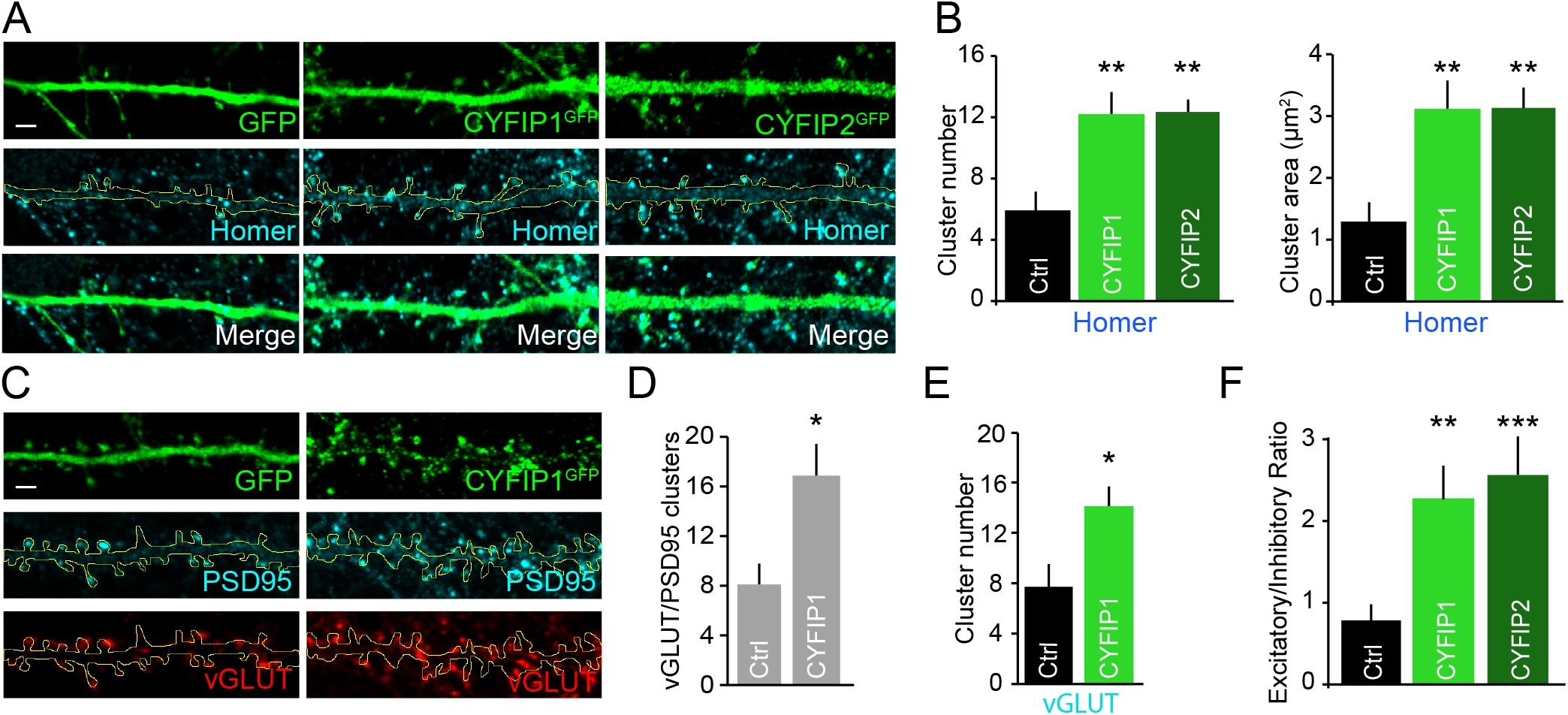
Increased expression of CYFIP1 and CYFIP2 alters the ratio of excitatory to inhibitory synapses. **(A)** Representative confocal images of hippocampal neurons transfected with CYFIP1^GFP^, CYFIP2^GFP^ or GFP control for 4 days before fixing at DIV14 and labelled with an antibody to the excitatory postsynaptic density protein homer. Scale bar, 2 μm. **(B)** Cluster analysis shows a significant increase in homer cluster number and area upon CYFIP1^GFP^ or CYFIP2^GFP^ overexpression (cluster number: from 5.9 ± 1.2 to 12.1 ± 1.5 for CYFIP1 and 12.3 ± 1.1 for CYFIP2; cluster area: from 1.3 ± 0.3 μm^2^ to 3.1 ± 0.5 μm^2^ for CYFIP1 and 3.1 ± 0.3 μm^2^ for CYFIP2; n = 17 cells from 3 preparations; Kruskal-Wallis oneway ANOVA, Dunn’s post-hoc multiple comparisons). **(C)** CYFIP1^GFP^ overexpressing hippocampal neurons labelled with antibodies to the excitatory pre and postsynaptic markers VGLUT and PSD95 respectively. Scale bar, 2 μm. **(D-E)** Cluster analysis revealed a significant increase in total number of excitatory synapses identified as VGLUT/PSD95 positive puncta (D) and VGLUT cluster number (E) upon CYFIP1^GFP^ overexpression (Total synapses: from 7 ± 1.6 to 13.6 ± 2; VGLUT number: from 7.7 ± 1.8 to 14 ±1.7; n = 15-16 cells from 3 preparations; p = 0.018 and 0.016; Student’s t-test). **(F)** The excitatory to inhibitory synaptic ratio quantified from neurons transfected with CYFIP1^GFP^, CYFIP2^GFP^ or GFP control and labelled with an antibodies to homer and GABA_A_R-γ2 as markers for excitatory and inhibitory synapses respectively (E/I ratio: from 0.8 ± 0.2 to 2.3 ± 0.4 for CYFIP1 and 2.6 ± 0.5 for CYFIP2; n = 17 cells from 3 preparations; Kruskal-Wallis one-way ANOVA, Dunn’s post-hoc multiple comparisons). *p < 0.05, **p < 0.01, ***p < 0.001. Bars indicate mean and error bars s.e.m.

Finally, we examined the ratio of inhibitory and excitatory synaptic clusters along dendrites in CYFIP1 or CYFIP2 overexpressing cells compared to control using antibodies against the GABA_A_R-γ2 subunit and homer, respectively. We observed a striking shift in the balance of excitatory and inhibitory synaptic puncta along dendrites upon CYFIP1/2 overexpression which led to a significant increase in the E/I ratio **(Fig. 3F**). Taken together, these results reveal that CYFIP protein overexpression differentially alters excitatory and inhibitory synapse number, disrupting the E/I synapse ratio.

### Disrupted inhibitory and excitatory synaptic activity in neurons overexpressing CYFIP1

Our results suggest that CYFIP1/2 overexpression has opposing effects on inhibitory and excitatory synapse integrity and hence, may dramatically impact neuronal excitation and inhibition. To further address this, we determined whether increased CYFIP1 dosage directly affects inhibitory and excitatory transmission in neurons, focusing on CYFIP1 as the gene has been more robustly associated with neurodevelopmental disorders. Whole-cell recordings were performed to measure inhibitory and excitatory transmission in neurons overexpressing CYFIP1 and co-expressing GFP **(Fig. S2**) (Kim et al., 2011). Analysis of mIPSCs from CYFIP1 overexpressing cells revealed a significant ~25 % decrease in mIPSC amplitude but no change in frequency compared to control neurons expressing GFP alone (**Fig. 4A-C**). The decreased mean mIPSC amplitude can be seen in the representative traces and in the leftward shift of the cumulative probability plot (**Fig 4D,E**). Overexpression of CYFIP1 had no effect on mIPSC kinetics (**Fig. 4F,G**). Conversely, when we analysed mEPSCs we observed no change in mEPSC amplitude but saw a robust and significant increase in mEPSC frequency (**Fig. 4H-J**). Again, this finding can be observed in both the example traces and the shift towards the right in the cumulative probability plot of mEPSC frequency (**Fig 4K,L**). The kinetics of mEPSCs were unchanged (**Fig. S3**). Finally, we measured the total charge transfer, a parameter that reflects both the amplitude and frequency of miniature synaptic events. The mean total charge transfer for mIPSCs was significantly decreased in CYFIP1 overexpressing cells while mEPSC charge transfer showed a trend towards an increase (**Fig. 4M,N**). These data demonstrate that CYFIP1 overexpression, not only alters synapse numbers, but results in functional deficits in synaptic transmission resulting in an imbalance of excitation and inhibition.

**Figure 4:**
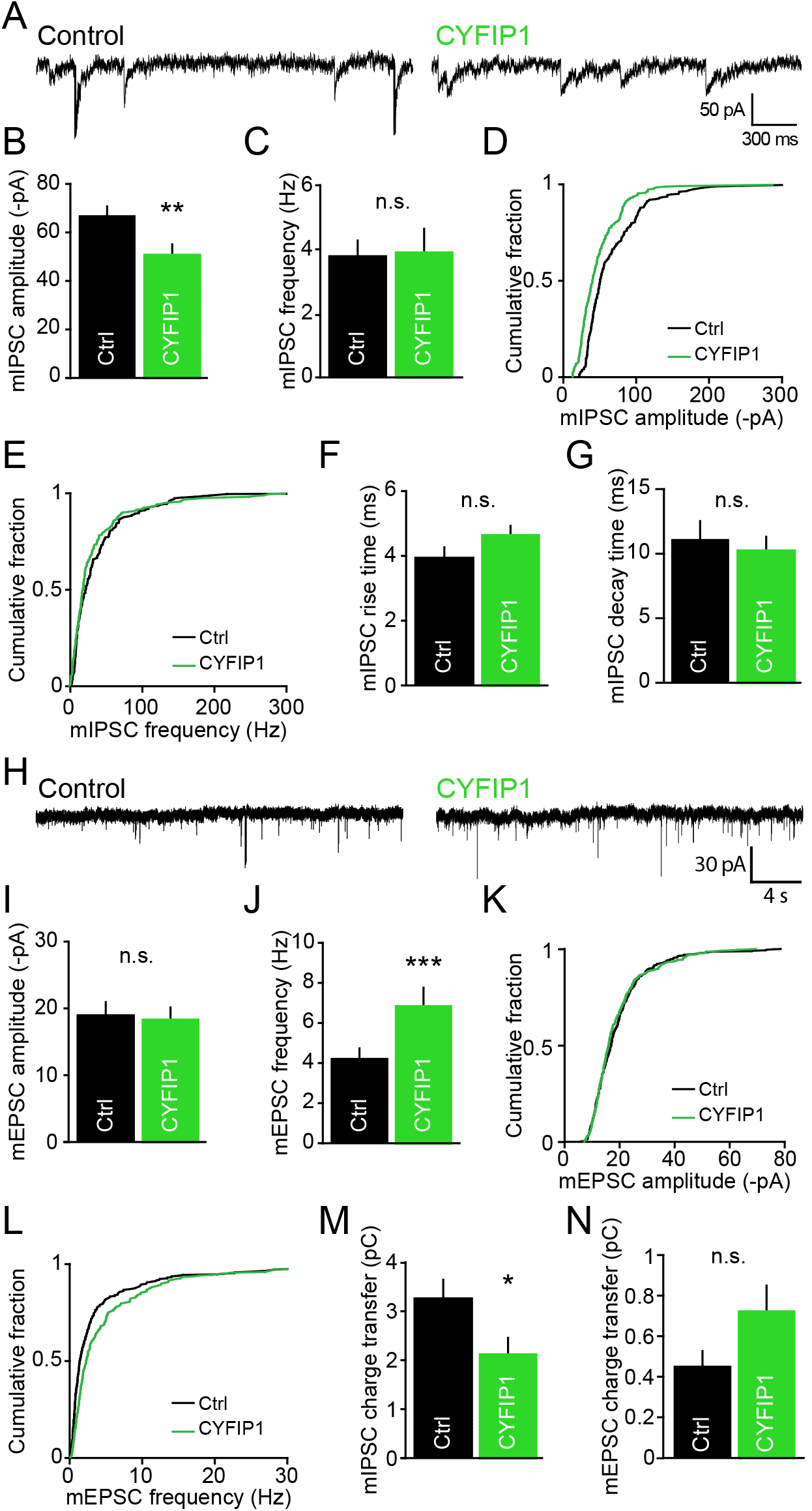
Increased CYFIP1 gene dosage disrupts inhibitory and excitatory synaptic transmission. **(A)** Representative traces of miniature inhibitory postsynaptic currents (mIPSCs) recorded from control GFP (Ctrl) and CYFIP1 overexpressing cultured hippocampal neurons at DIV14-16. **(B-C)** Pooled data of mIPSCs showing neurons transfected with CYFIP1 have a reduction in (B) mean mIPSC amplitude but no change in (C) mean mIPSC frequency (mIPSC amplitude: from 66.9 ± 3.8 -pA to 50.7 ± 3.6 -pA, p = 0.0062; frequency: from 3.8 ± 0.5 Hz to 3.9 ± 0.7 Hz, p = 0.92 n.s.; all n = 10 cells from 3 preparations; Student’s t-test). **(D-E)** Cumulative frequency graphs of mIPSC (D) amplitude and (E) frequency. **(F)** Graph of mIPSC rise time kinetics (from 4 ± 0.3 ms to 4.6 ± 0.3 ms; n = 9 cells from 3 preparations; p = 0.0623 n.s.; Mann-Whitney). **(G)** Graph of mIPSC decay time kinetics (from 11.1 ± 1.4 ms to 10.2 ± 1.1 ms; n = 10-11 cells from 3 preparations; p = 0.618 n.s.; Student’s t-test). **(H)** Representative traces of miniature excitatory postsynaptic currents (mEPSCs) recorded from CYFIP1 or GFP control (Ctrl) transfected neurons. **(I-J)** Pooled data of mEPSCs showing neurons transfected with CYFIP1 have no difference in (I) mean mEPSC amplitude but a significant increase in (J) mean mEPSC frequency compared with control (mEPSC amplitude: from 19.0 ± 1.7 -pA to 17.0 ± 1.4 -pA, p = 0.367 n.s.; frequency: from 1.5 ± 0.1 Hz to 2.9 ± 0.4 Hz, p = 0.0003; n =14-120 cells from 3 preparations; Student’s t-test). **(K-L)** Cumulative frequency graphs of mEPSC (K) amplitude and (L) frequency. **(M)** Quantification of mIPSC total charge transfer demonstrating that CYFIP1 overexpression causes a significant decrease in mean charge transfer compared to GFP control expressing cells (from 3.3 ± 0.4 pC to 2.1 ± 0.3 pC; n = 9 cells from 3 preparations; p = 0.0341; Student’s t-test). **(N)** Quantification of mEPSC total charge transfer demonstrating that CYFIP1 overexpression causes a trend towards an increase in mean charge transfer compared to GFP expressing control cells (mEPSC charge transfer: from 0.45 ± 0.1 pC to 0.7 ± 0.1 pC; n = 10-11 cells from 3 preparations; p = 0.1307 n.s.; Mann-Whitney). *p < 0.05, **p < 0.01, ***p < 0.001. Bars indicate mean and error bars s.e.m.

### Decreased CYFIP1 gene dosage alters neuronal and dendritic spine morphology

*CYFIP1* can undergo microdeletion as well as microduplication, therefore, it was equally important to study the effect of loss of CYFIP1 on synaptic inhibition and the E/I balance. However, as previously described, the CYFIP1 constitutive knockout (KO) mouse is embryonic lethal and hence, the impact of deleting all CYFIP1 in the brain remains undetermined (Pathania et al., 2014). To circumvent embryonic lethality and study cell type specific effects of CYFIP1 deletion we generated a conditional KO (cKO) mouse line selectively lacking CYFIP1 in forebrain excitatory neurons using a Nex-Cre driver line (Goebbels et al., 2006; Skarnes et al., 2011; White et al., 2013). In this Cre driver line, Cre activity was observed in neocortex and hippocampus from around embryonic day 12 (E12) onwards. This allowed us to determine the impact of deleting CYFIP1 through development specifically in excitatory cells from these brain regions (**Fig. 5A and SFig. 3A**). CYFIP1^NEX^ cKO animals were viable until adulthood with no obvious abnormalities. Western blotting of post-natal day (P) 30 hippocampal brain lysates with a CYFIP1 specific antibody revealed a robust reduction of CYFIP1 expression in CYFIP1^NEX^ cKO mice compared to control floxed animals (**Fig. 5B,C**). Remaining CYFIP1 expression detected in western blots presumably comes from CYFIP1 in other cell populations such as interneurons and glia. Fluoronissl labelling of thin brain sections revealed that CYFIP1^NEX^ cKO mice did not show any gross morphological abnormalities in neocortical and hippocampal brain structure when compared to control (**Fig. 5D**).

**Figure 5:**
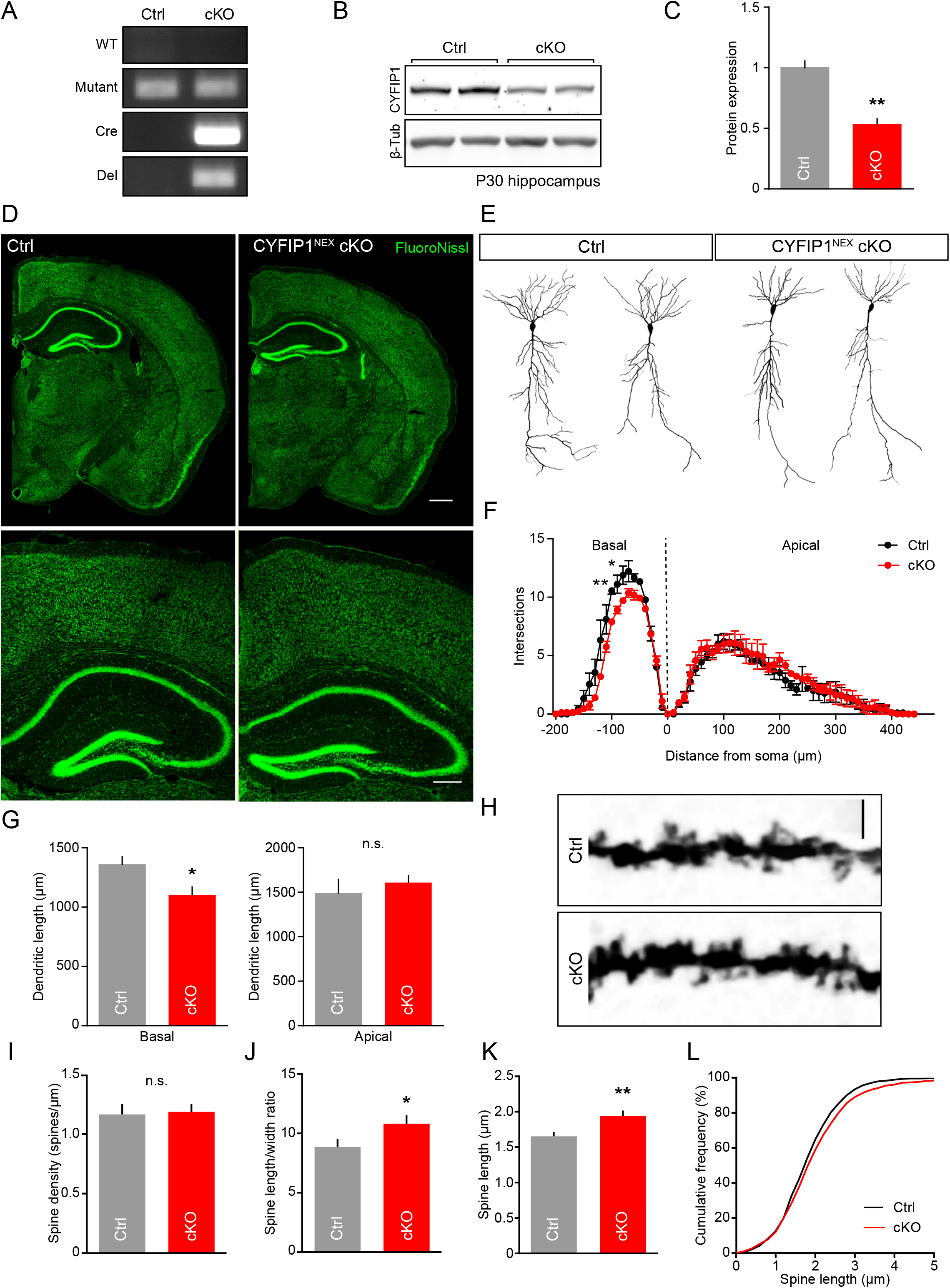
Loss of CYFIP1 expression in principal cells of the neocortex alters hippocampal cell morphology. **(A)** PCR genotyping of CYFIP1^NEX^ conditional knockout (cKO) animals generated from the KO-first strategy. CYFIP1 floxed animals were crossed with mice expressing cre recombinase under the Nex promoter. Animals were genotyped with wild type (WT), mutant, cre recombinase (Cre) and deletion (Del) primers. See Figure S3 for details. **(B,C)** Western blot analysis and quantification displaying fold change of CYFIP1 protein levels from floxed control (Ctrl) and CYFIP1^NEX^ conditional KO (cKO) P30 hippocampal brain lysates (from 1 ± 0.1 to 0.5 ± 0.05, n = 3 animals per condition, p = 0.0033, Student’s t-test). **(D)** FlouroNissl staining of control floxed (Ctrl) and CYFIP1^NEX^ cKO P30 mouse coronal brain sections shows no major abnormalities in gross brain morphology between the two genotypes. Scale bar, 500 μm, zoom 250 μm. **(E)** Example reconstructions of CYFIP1^NEX^ cKO and floxed littermate control (Ctrl) Golgi-stained P30 CA1 neurons. **(F)** Sholl analysis of CYFIP1^NEX^ cKO (cKO) CA1 neurons compared to control cells (Ctrl) to measure dendritic complexity. CYFIP1^NEX^ cKO basal dendrites are less complex (Basal dendrites −100 μm: p < 0.05; −120 μm: p < 0.01; 2-way ANOVA, Bonferroni’s multiple comparisons). **(G)** Total dendritic length of CYFIP1^NEX^ cKO (cKO) CA1 neurons compared to control cells (Ctrl). CYFIP1^NEX^ cKO basal dendrites have significantly less dendritic length (Dendritic length: basal, from 1360 ± 65.9 μm to 1099 ± 70.5 μm, p = 0.0178; apical, from 1490 ± 153.5 μm to 1609 ± 78 μm, p = 0.46 n.s.; n = 9-13 reconstructed cells from 3 animals per genotype, Student’s t-test). **(H)** Example dendrites and dendritic spines of CYFIP1^NEX^ cKO (cKO) and floxed littermate control (Ctrl) Golgi-stained P30 CA1 neurons. Scale bar, 5 μm. **(I-K)** Dendritic spine analysis revealed no change in spine density (I) between CYFIP1^NEX^ cKO (cKO) and floxed littermate control (Ctrl) neurons but an increase in the spine length:width ratio (J) in CYFIP1^NEX^ cKO neurons as a result of an increase in dendritic spine length (K) (Spine density: from 1.2 ± 0.1 spines/μm to 1.2 ± 0.1 spines/μm, p = 0.32 n.s.; length:width ratio: from 8.9 ± 0.7 to 10.8 ± 0.7, p = 0.0407; length: from 1.7 ± 0.1 μm to 1.9 ± 0.1 μm, p = 0.02; n = 45 dendritic processes from 3 animals per genotype, Student’s t-test). **(L)** Cumulative frequency graph of dendritic spine length. *p < 0.05, **p < 0.01. Bars indicate mean and error bars s.e.m.

CYFIP1 haploinsufficiency in constitutive CYFIP1 heterozygous KO mice led to decreased dendritic complexity and altered dendritic spine maturation both *in vitro* and *in vivo* (Pathania et al., 2014). Therefore, we initially assessed dendritic morphology in hippocampal neurons from CYFIP1^NEX^ cKO mice. Golgi-stained CA1 neurons analysed from P30 CYFIP1^NEX^ cKO brains showed significantly less dendritic complexity in the basal compartment compared to neurons analysed from littermate control tissue. Consistent with this, total basal dendritic length was reduced however, branch point number was unchanged (**Fig. 5E-G**). The impact on dendritic spines of completely knocking out CYFIP1 in principal cells was unexpectedly subtle. Spine density was unchanged in CYFIP1^NEX^ cKO neurons. There was however, a significant increase in the spine length to width ratio, as a result of a significant increase in spine length, consistent with the spine phenotypes previously reported upon constitutive CYFIP1 heterozygous KO (Pathania et al., 2014) (**Fig. 5H-L**). Taken together, these data illustrate that the CYFIP1^NEX^ cKO mice show similar deficits in dendrite morphology and spine maturation to those that have been described for a CYFIP1 haploinsufficient model and support these effects to be cell autonomous to the principal cells.

### Postsynaptic loss of CYFIP1 increases inhibitory synapse size and strength

To further explore the impact of CYFIP1 deletion on synaptic components we probed hippocampal lysates from P30 control and CYFIP1^NEX^ cKO brains with antibodies to key molecular components of the inhibitory and excitatory PSDs. Interestingly, while the levels of key excitatory postsynaptic proteins including homer and PSD95 were unchanged we observed a significant increase in the levels of the inhibitory GABA_A_R-β2/3 subunits and the ASD-associated neuroligin 3 adhesion molecule, which can be found at both inhibitory and excitatory postsynapses (**Fig. 6A**). CYFIP1 loss of function may therefore have an opposite effect to that of upregulation, causing an increase in inhibitory synapse stability. To validate this we carried out immunohistochemistry on thin hippocampal sections taken from P30 control floxed and CYFIP1^NEX^ cKO brains. Sections were labelled with antibodies to VGAT and gephyrin to report inhibitory pre and postsynapses and DAPI to indicate cell bodies. Quantification in the *stratum pyramidale* layer of the hippocampus revealed a significant increase in gephyrin cluster area in cKO tissue compared to control while VGAT cluster area was unchanged (**Fig. 6B-D**). These data highlight that loss of CYFIP1 *in vivo* in glutamatergic principal cells results in an increase in inhibitory synapse size and the levels of inhibitory synaptic proteins.

**Figure 6:**
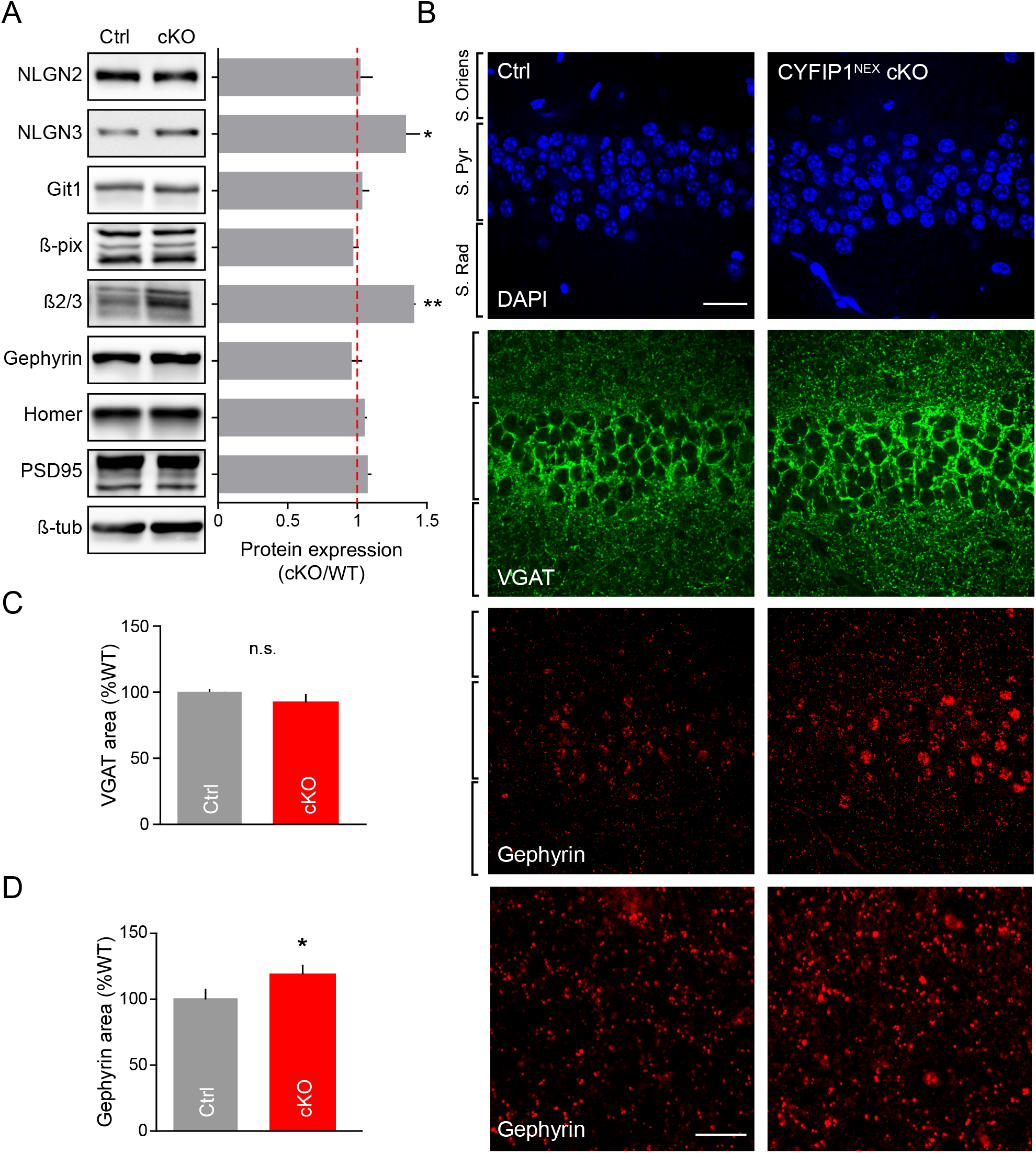
Decreased CYFIP1 gene dosage alters expression of inhibitory scaffold molecules and inhibitory synaptic structure *in vivo*. **(A)** Western blot analysis and quantification displaying protein expression ratios of inhibitory and excitatory postsynaptic proteins from control (Ctrl) and CYFIP1^NEX^ conditional KO (cKO) P30 hippocampal brain lysates (neuroligin 2 (NLGN2): 1.02 ± 0.09; neuroligin 3 (NLGN3): 1.35 ± 0.1, p = 0.0286; Git1: 1.03 ± 0.05; β-pix: 0.97 ± 0.04; GABA_A_R β2/3: 1.41 ± 0.01, p = 0.0017; gephyrin: 0.96 ± 0.07; homer: 1.05 ± 0.02; PSD95: 1.07 ± 0.03; n = 3 animals per condition; Student’s t-test). **(B)** Confocal images of adult control (Ctrl) and CYFIP1^NEX^ cKO hippocampal brain sections immunolabelled with antibodies to VGAT and gephyrin, co-stained with DAPI. Scale bar, 25 μm, zoom 10 μm. **(C,D)** Normalised total cluster area quantification of CYFIP1^NEX^ cKO (cKO) tissue as a percentage of floxed control (Ctrl) showing a no change in (C) VGAT cluster area and an increase in (D) gephyrin cluster area (VGAT: from 100 ± 2.4 % to 94 ± 5.2 %, p = 0.336 n.s.; gephyrin: from 100 ± 6.9 % to 119 ± 6.1 %, p = 0.0473; n = 18 hippocampal regions from 3 animals per genotype; Student’s t-test) in CYFIP1 cKO tissue compared to control. *p < 0.05, **p < 0.01. Bars indicate mean and error bars s.e.m.

Finally, we investigated whether the changes in inhibitory synapses observed in CYFIP1^NEX^ cKO mice translated into a functional effect on synaptic transmission. We examined mIPSCs in acute hippocampal slices from control and CYFIP1^NEX^ cKO P28-34 mice, in which CA1 pyramidal cells could be identified unambiguously. Recordings from these cells showed that deletion of CYFIP1 resulted in a significant increase in mIPSC amplitude consistent with a shift to the right in the cumulative frequency plot of mIPSC amplitude. No change was observed in mIPSC frequency and mIPSC rise and decay time between control and CYFIP1^NEX^ cKO neurons (**Fig. 7A-D**). Importantly, CYFIP1 deletion had no effect on AMPAR-mediated mEPSCs (**Fig. 7E-H**), confirming a selective effect on synaptic inhibition in P28-34 animals. Lastly, we measured the total charge transfer mediated by both inhibitory and excitatory postsynaptic currents. This showed that mIPSC charge transfer was increased by ~70% in CYFIP1 deleted cells compared to control while mEPSC charge transfer was unchanged (**Fig. 7I**). The probability curve of mIPSC and mEPSC charge transfer from CYFIP1^NEX^ cKO neurons normalised to control demonstrates the resultant imbalance between inhibitory and excitatory transmission observed with loss of CYFIP1 expression (**Fig. 7J**). Thus, CYFIP1 deletion appears to have a dramatic impact on inhibitory synapse integrity and the strength of inhibition.

**Figure 7:**
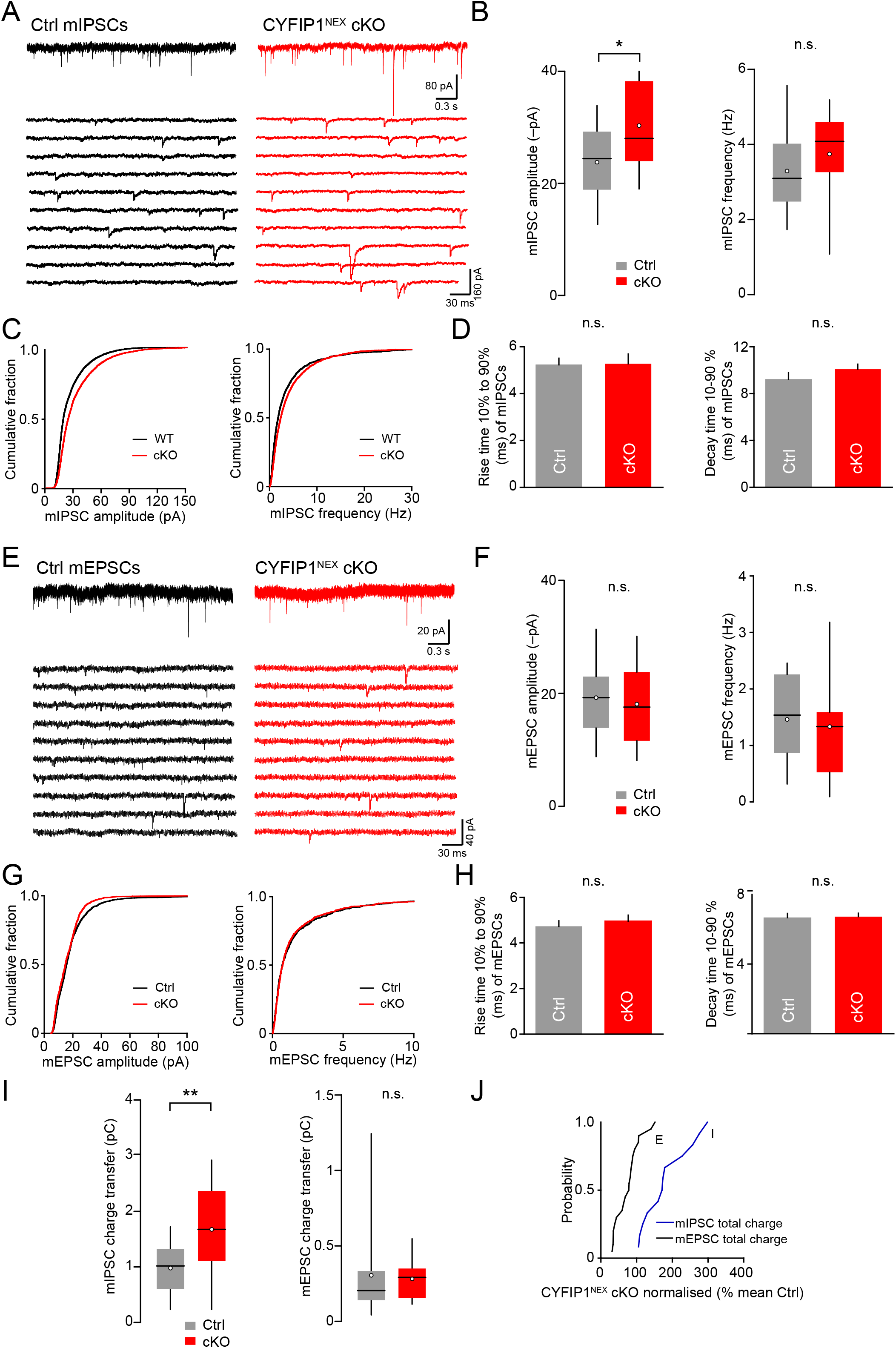
Postsynaptic loss of CYFIP1 *in vivo* increases inhibitory synaptic function. **(A)** Representative recordings of miniature inhibitory postsynaptic currents (mIPSCs) (-70 mV) in CA1 pyramidal cells from P28-34 control floxed (left) and CYFIP1^NEX^ cKO mice (right). Lower panels are representative sections of recordings (contiguous 0.3 s segments). **(B)** Pooled data showing increase mIPSC mean amplitude in CYFIP1^NEX^ cKO mice (cKO) (from 23.8 ± 1.8 -pA to 30.3 ± 2.2 -pA, n = 13-14 cells, p = 0.0288) but lack of change in frequency (from 3.3 ± 0.3 Hz to 3.7 ± 0.3 Hz, n = 13-15 cells, p = 0.324 n.s.) All Student’s t-test. Box-and-whisker plots indicate median (line), 25-75th percentiles (box), range of data within 1.5 x IQR of box (whiskers) and mean (open circles). **(C)** Cumulative frequency graphs of mIPSC amplitude (left) and frequency (right). **(D)** Graphs of mIPSC kinetics showing no change in rise and decay time between control (Ctrl) and CYFIP1^NEX^ cKO mice (cKO) (rise time: from 5.2 ± 0.3 ms to 5.3 ± 0.4 ms, n = 13-14 cells, p = 0.935 n.s.; decay time: 9.2 ± 0.6 ms to 10.1 ± 0.4, n = 13-15 cells, p = 0.247 n.s., both Student’s t-test). **(E)** Representative recordings of miniature excitatory postsynaptic currents (mEPSCs) (-70 mV) in CA1 pyramidal cells from P28-34 control floxed (left) and CYFIP1^NEX^ cKO mice (right). Lower panels are representative sections of recordings (contiguous 0.3 s segments). **(F)** Pooled data showing a lack of change in mEPSC mean amplitude (from 19.2 ± 1.4 -pA to 18 ± 1.4 -pA, n = 21-22 cells, p = 0.549 n.s.) and frequency between control (Ctrl) and CYFIP1 cKO mice (cKO) (from 1.5 ± 0.2 Hz to 1.3 ± 0.2 Hz, n = 18-20 cells, p = 0.565 n.s.). All Student’s t-test. Box-and-whisker plots indicate median (line), 25-75th percentiles (box), range of data within 1.5 x IQR of box (whiskers) and mean (open circles). **(G)** Cumulative frequency graphs of mEPSC amplitude (left) and frequency (right). **(H)** Graphs of mEPSC kinetics showing no change in rise and decay time between control (Ctrl) and CYFIP1^NEX^ cKO mice (cKO) (rise time: from 4.7 ± 0.2 ms to 5 ± 0.3 ms; n = 22 cells, p = 0.527 n.s.; decay time: from 6.7 ± 0.2 ms to 6.7 ± 0.2 ms; n = 22 cells, p = 0.8415 n.s.; both Student’s t-test). **(I)** Pooled data showing increase in mIPSC mean charge transfer in CYFIP1^NEX^ cKO (cKO) mice compared to floxed control (Ctrl) (from 1 ± 0.2 pC to 1.7 ± 0.2 pC, n = 13-14 cells, p = 0.0067) but no change in mEPSC mean charge transfer (from 0.3 ± 0.1 pC to 0.3 ± 0.03 pC, n = 18-20 cells, p = 0.753 n.s.) All Student’s t-test. Box-and-whisker plots indicate median (line), 25-75th percentiles (box), range of data within 1.5 x IQR of box (whiskers) and mean (open circles). **(J)** The probability curve of mean mEPSC and mIPSC charge transfer in CYFIP1^NEX^ cKO mice as a percentage of control mice highlighting the imbalance between excitation and inhibition in CYFIP1^NEX^ cKO animals. *p < 0.05, **p < 0.01. Bar graph bars indicate mean and error bars s.e.m.

## Discussion

Alterations in E/I balance are implicated in neuropsychiatric disorders including ASD and SCZ (Foss-Feig et al., 2017) but how this may be caused by genetic variation is poorly understood. Here we report that the ASD and SCZ associated protein CYFIP1 is localised to inhibitory synapses and can regulate inhibitory synapse stability and the balance between neuronal excitation and inhibition. CYFIP1 upregulation (modelling CNV gain) resulted in an increase in the excitatory to inhibitory synaptic ratio consistent with a functional decrease in mIPSC amplitude and increase in mEPSC frequency. In contrast, CYFIP1 loss had the opposite effect on neuronal inhibition, leading to increased inhibitory postsynaptic clustering, enhanced expression of neuroligin 3 and GABA_A_R β-subunits and an increase in mIPSC amplitude in CA1 hippocampal cells. Our data provides strong support for altered inhibition and disruption in the E/I balance being a pathological consequence of *CYFIP1* CNV and points towards disruption in inhibitory synaptic structure and function as being part of the underlying mechanism.

CYFIP1 was previously shown to be enriched at excitatory synapses where it can regulate F-actin dynamics (Pathania et al., 2014; De Rubeis et al., 2013), protein translation (Napoli et al., 2008) and dendritic spine structural plasticity (Pathania et al., 2014). Here we now demonstrate that CYFIP1 and CYFIP2 are also enriched at inhibitory synapses where they colocalise with the inhibitory postsynaptic scaffold gephyrin opposed to VGAT positive presynaptic terminals. CYFIP1 was also found to interact with gephyrin, supporting its intimate association with the inhibitory postsynaptic domain and suggesting it may also be found in complexes with GABA_A_Rs. Moreover, STED imaging revealed CYFIP1 and gephyrin to be in closely adjacent clusters consistent with the sub-synaptic localisation of other inhibitory synaptic enriched proteins (Woo et al., 2013). Interestingly, we further demonstrate that increased CYFIP1 dosage leads to a decrease in the size of inhibitory synapses and reduced synaptic inhibition likely due to a loss of surface γ2-subunit containing synaptic GABA_A_R clusters.

Intriguingly, increased CYFIP1 dosage has the opposite effect on excitatory synapses, leading to increased VGLUT positive presynaptic and homer positive postsynaptic cluster number and area and thus increased excitatory synapse number. In particular, a high proportion of these synapses appeared on the dendritic shaft compared to spines, perhaps due to the molecular and spatial confinement present within the spines as we detected significantly more long thin spines on CYFIP1 overexpressing cells. This subtype of spines is proposed to reflect a more immature spine structure and contain fewer and less established synapses and has been repeatedly identified in rodent models and patients with ASD (Phillips and Pozzo-Miller, 2015). In addition to an increase in the number of thin spines we also detected more mushroom spines, which may reflect the increase in the number of established synapses observed.

An increase in dendritic spine density and number of aberrant spines was recently observed in a transgenic mouse which overexpresses *Cyfip1* under the endogenous promoter (Oguro-Ando et al., 2014). Our own results also demonstrate an increase in synapse number and altered spine morphologies correlated with a marked increase in mEPSC frequency following CYFIP1 overexpression. Given that CYFIP1 is only upregulated in the postsynaptic compartment due to sparsely transfected neurons in this experiment (i.e. most inputs will be from non-transfected cells), this frequency increase likely reflects a postsynaptically driven increase in excitatory synapse number, rather than any presynaptic effect on release probability (Hsiao et al., 2016). Thus along with decreased inhibition, upregulation of CYFIP1 expression also leads to increased excitation and together these two effects would likely lead to altered E/I balance.

Upregulation of CYFIP2 phenocopies both the inhibitory and the excitatory alterations in synapse number and size and the spine morphology changes observed with increased CYFIP1 dosage. Although CYFIP2 CNVs have yet to be reported, alterations in CYFIP2 function have been associated with neurological and neuropsychiatric disorders (Han et al., 2015; Kumar et al., 2013; Nakashima et al., 2018; Tiwari et al., 2016). Moreover, CYFIP2 protein levels have been found to be increased in brain tissue from patients with SCZ and fragile X syndrome and decreased in Alzheimer’s disease (Föcking et al., 2014; Hoeffer et al., 2012; Tiwari et al., 2016) while CYFIP2 happloinsufficiency or mutations in mice led to altered dendritic spine morphology in the cortex (Han et al., 2015; Kumar et al., 2013; Tiwari et al., 2016) and autism-like behaviours (Han et al., 2015). Our data suggests that, in addition to spine defects, disruptions in CYFIP2 expression lead to altered synaptic inhibition, resulting from defects in synapse structure and the E/I synapse ratio, which may contribute to the molecular mechanisms underlying CYFIP2-associated neurological disorders.

In addition to studying the impact of CYFIP1 upregulation we also determined the impact of CYFIP1 loss on neuronal development and connectivity. Constitutive KO of CYFIP1 leads to early embryonic lethality which has thus far limited CYFIP1 loss of function studies to exploring the impact of CYFIP1 haploinsufficiency in the heterozygous KO model (Bozdagi et al., 2012; Chung et al., 2015; Hsiao et al., 2016; Pathania et al., 2014; De Rubeis et al., 2013). While these have been informative, given the strong genetic links of CYFIP1 to neurological disease it is clearly also important to establish the impact of complete loss of CYFIP1 in CNS neurons and the extent to which the effects of CYFIP1 dysfunction are cell autonomous to glutamatergic neurons where most studies have focused. To this end, we developed a CYFIP1^NEX^ cKO where CYFIP1 was deleted from principal cells of the neocortex. Surprisingly, CYFIP1^NEX^ cKO resulted in relatively mild defects in dendritic branching and spine maturation in P30 hippocampal principal cells, quite similar to those previously reported upon CYFIP1 haploinsufficiency (Pathania et al., 2014); perhaps due to compensation from CYFIP2 (Han et al., 2015; Pathania et al., 2014). Indeed, CYFIP1^NEX^ cKO animals did not exhibit alterations in excitatory synaptic transmission at this age consistent with reports of adult CYFIP1 constitutive heterozygous KO mice where basal excitatory synaptic transmission was unaltered (Bozdagi et al., 2012; Hsiao et al., 2016). Importantly, the cell selectivity of our newly reported CYFIP1^NEX^ cKO supports dendritic branching and spine alterations upon disrupted CYFIP1 expression (Oguro-Ando et al., 2014; Pathania et al., 2014; De Rubeis et al., 2013) to be primarily cell autonomous to the principal cells.

While hippocampal expression levels of key excitatory synapse components including homer and PSD95 were unaffected by conditional CYFIP1 deletion, we found increased expression levels of GABA_A_R-β2/3 subunits supporting a greater impact on inhibitory synapse function. The majority of synaptic GABA_A_Rs require the incorporation of β2/3 subunits to form and be efficiently trafficked to the plasma membrane where they can be tethered by gephyrin (Luscher et al., 2011). Therefore, higher levels of GABA_A_R β2/3 subunits likely reflect increased numbers of synaptic GABA_A_Rs. In line with this, we observed an increase in synaptic gephyrin clustering and an increase in mIPSC amplitude in the hippocampus upon conditional CYFIP1 loss from principal cells. Thus at this age the main effect of CYFIP1 deletion appears to be an increase in the strength of synaptic inhibition.

Intriguingly, CYFIP1^NEX^ cKO also led to a significant increase in hippocampal neuroligin 3 (NLGN3) protein levels. Neuroligin 3 is a member of the neuroligin 1-4 family of synaptic adhesion molecules, which form trans-synaptic interactions to drive synapse specification and maintenance and have been genetically implicated in neurodevelopmental disorders (Bemben et al., 2015a, 2015b; Jamain et al., 2003). Neuroligin 1 and neuroligin 2 are exclusively localised to excitatory and inhibitory synapses, respectively (Poulopoulos et al., 2009; Varoqueaux et al., 2004). In contrast, neuroligin 3 can be found at both types of synapse and at inhibitory synapses can interact with gephyrin and neuroligin 2 (Budreck and Scheiffele, 2007). It is intriguing that neuroligin 3 is found to be upregulated upon CYFIP1 deletion while neuroligin 2 expression levels were unaltered and equally surprising that its upregulation appears to only enhance synaptic inhibition (but not excitation). However, our findings are consistent with recent work demonstrating that overexpression of neuroligin 3 leads to a selective increase in the strength of inhibition over excitation (Chanda et al., 2017; Fekete et al., 2015). By stabilising GABA_A_Rs at synapses, upregulated neuroligin 3 may contribute to the increased levels of GABA_A_R-β2/3 subunits observed and explain mechanistically the selective increase in synaptic inhibition. Thus, neuroligin 3 expression levels may play a pivotal role in bi-directional control of the excitatory to inhibitory synapse ratio and the E/I balance. Indeed ASD-associated point mutations in NLGN3 have been show to impact the E/I synaptic ratio in neurons (Tabuchi et al., 2007; Zhang et al., 2016) and NLGN3 KO mice display ASD behavioural deficits such as reduced ultrasound vocalisation and deficits in social novelty preference (Radyushkin et al., 2009).

The mechanisms by which CYFIP1 expression levels can bi-directionally regulate postsynaptic inhibition remain to be fully elucidated. CYFIP1 in the WRC can regulate actin dynamics by complexing with the actin regulator Rac1 (Chen et al., 2010; Kobayashi et al., 1998) and altered CYFIP1 expression levels may disturb inhibition not only through altered ARP2/3 activity, but also by perturbing the balance of Rac1 signalling - itself a key regulator of GABA_A_R synaptic stabilisation and synaptic inhibition (Smith et al., 2014). Altered actin dynamics through WRC and Rac1 signalling also likely account for the impact of CYFIP1 levels on dendritic spine dynamics as previously proposed (Pathania et al., 2014; De Rubeis et al., 2013). Interestingly, neuroligin 3 (but not neuroligin 2) contains a newly identified WIRS (<u>W</u>RC <u>I</u>nteracting <u>R</u>eceptor <u>S</u>equence) peptide motif that binds to a key protein-binding pocket between CYFIP1 and Abi unique to the intact, fully assembled WRC (Chen et al., 2014; Chia et al., 2014). Disrupted coupling of neuroligin 3 to the WRC upon CYFIP1 deletion might alter its trafficking, by disrupting its surface downmodulation or intracellular sorting (Anitei et al., 2010; Xu et al., 2016), leading to increased surface levels and reduced degradation. Altered surface stability or turnover of neuroligin 3 might work hand in hand with increased neuroligin 3 expression due to relief of FMRP-dependent translational repression upon CYFIP1 loss (Darnell et al., 2011; Napoli et al., 2008).

E/I balance shift can lead to deficits in network activity, disrupted information processing and altered behaviours (Blundell et al., 2009; Crestani et al., 1999; Gkogkas et al., 2013; Tora et al., 2017; Yizhar et al., 2011). An increase in the ratio of excitatory to inhibitory synapses as observed upon CYFIP1 upregulation is consistent with altered E/I balance in ASD and in mouse models of numerous neuropsychiatric disorders (Bateup et al., 2013; Chao et al., 2010; Gao and Penzes, 2015; Nelson and Valakh, 2015; Paluszkiewicz et al., 2011; Smith et al., 2017) and the increased risk of neuropsychiatric symptoms in some individuals with *CYFIP1* duplication. Intriguingly, patients with temporal lobe epilepsy and pilocarpine-treated rats show an upregulation of CYFIP1 expression consistent with the notion that increased CYFIP1 expression is associated with altered E/I balance and associated behaviours (Huang, 2015). Enhanced inhibition upon CYFIP1 deletion could contribute to the intellectual disability and cognitive changes reported in individuals with 15q11.2 microdeletions. Indeed, excess inhibition contributes to cognitive impairment in Down’s syndrome models where disrupted long term potentiation, learning and memory can be improved by pharmacologically targeting GABA_A_Rs (Rudolph and Möhler, 2014). Interestingly, overexpression of neuroligin 3 in hippocampal principal cells was also recently reported to selectively increase synaptic inhibition by somatostatin expressing interneurons that innervate distal dendrites at the expense of perisomatic inputs from parvalbumin expressing interneurons (Horn and Nicoll, 2018). Whether CYFIP1 deletion and concomitant neuroligin 3 upregulation could similarly alter the balance of inhibitory circuit control by these two types of interneuron remains to be determined.

Our results have established a link between altered CYFIP1 dosage, changes in synaptic inhibition and excitation, and altered E/I balance. This provides important new insights into the role CYFIP proteins have in synaptic function and network activity and how *CYFIP1* dysregulation in 15q11.2 CNV may impact CNS function to contribute to the development of neuropsychiatric and neurodevelopmental disorders. Furthermore, our work supports the idea that synaptic inhibition is a therapeutic target and that drugs acting on GABA_A_Rs may prove beneficial for individuals harbouring CYFIP1 CNVs.

## Methods

Details regarding animals, antibodies, cDNA cloning, primary neuronal culture, preparation of brain lysates, PLA procedures and Golgi staining are included in the Supplemental Experimental Procedures.

### Immunocytochemistry and immunohistochemistry

Hippocampal cultures were fixed in 4 % PFA (PBS, 4 % paraformaldehdye, 4 % sucrose, pH 7) for 7 minutes then permeablised for 10 minutes in block solution (PBS, 10 % horse serum, 0.5 % BSA, 0.2 % Triton X-100). Coverslips were incubated with primary antibody diluted in block solution for 1 hour, washed in PBS, then incubated for 1 hour with secondary antibody. Finally coverslips were washed and mounted onto glass slides using ProLong Gold antifade reagent (Invitrogen). For surface labelling, block solution was used without detergent. The in situ proximity ligation assay (PLA) was used according to the manufacturer’s instructions (Duolink^®^ PLA technology, SIGMA, see Supplemental Experimental Procedures).

For immunohistochemistry, adult mouse brains of either sex were fixed in 4 % PFA overnight and cryoprotected in 30 % sucrose/PBS solution overnight before freezing at −80 °C. The brain samples were embedded in tissue freezing compound and 30 μm brain sections were generated using a Cryostat (Bright Instruments, Luton, UK). Free floating thin sections were permeablised for 4-5 hours in block solution (PBS, 10 % horse serum, 0.5 % BSA, 0.5 % Triton X-100, 0.2 M glycine) then incubated with primary antibody diluted in block solution overnight at 4 °C. For mouse primary antibodies, slices were first incubated overnight at 4 °C with mouse Fab fragment (1:50 with block solution; 115-007-003, Jackson ImmunoResearch, West Grove, PA, USA) to reduce background staining on the mouse tissue. Slices were washed 4-5 times in PBS for 2 hours then incubated for 3-4 hours with secondary antibody at room temperature. The slices were then washed 4-5 times in PBS for 2 hours and mounted onto glass slides using Mowiol mounting medium. For antigen retrieval, slices were incubated in sodium citrate solution at 80°C for 40 mins and then washed 3x in PBS prior to blocking.

### Confocal microscopy and image analysis

All confocal images were acquired on a Zeiss LSM700 upright confocal microscope using a 63X oil objective (NA: 1.4) unless otherwise stated. For synaptic localisation, enrichment and cluster analysis experiments from cultured neurons, a single plane image of each cell was captured using a 0.5X zoom. From this, 3 sections of primary or secondary dendrite, ~100μm from the soma, were imaged with a 3.5X zoom (equating to a 30μm length of dendrite). For brain sections from adult male and female fixed brains, 2 low magnification regions of the hippocampus were captured using a 63X objective and 0.5X zoom. From this, 3 regions were imaged within each hippocampal strata with a 2X zoom for analysis. Acquisition settings and laser power were kept constant within all experiments.

Line scans used for protein localisation were performed in ImageJ using the PlotProfile function (NIH, Bethesda, MD, USA), pixel intensity was calculated as a function of distance along a manually drawn line and plotted on a graph. Synaptic enrichment and cluster analysis was carried out using Metamorph (Molecular Devices, Sunnyvale, CA, USA). Analysis was carried out on the zoom dendrite images and then averaged to give a value per cell. To quantify protein enrichment at synaptic sites, the protein fluorescence intensity was measured as the average intensity within the labelled synaptic puncta and normalised to the average intensity of the total process. For synaptic cluster analysis, the length of dendrite was traced to generate a dendritic region of interest (ROI). This ROI was transferred to all cluster channels. A user-defined threshold was then applied to each synaptic marker channel and regions were generated around the thresholded area within the dendrite ROI. Number of regions and total area of regions per 30μm of dendrite were quantified as a readout for synaptic clusters. Clusters smaller than 0.01μm^2^ were excluded from the number of regions analysis. Thresholds were set individually for each cluster channel and kept constant across treatment conditions within an experiment. For brain sections labeled with antibodies against gephyrin and VGAT, the Synapse Counter plugin for ImageJ (NIH, Bethesda, MD, USA) was used. Background subtraction and max filter parameters were set to 10 and 1 respectively. Clusters greater than 0.095 μm^2^ and less than 1 μm^2^ were considered for total cluster area analysis. For spine morphology analysis of cultured neurons, confocal image stacks were acquired. Spines were manually identified on 100-200μm long dendritic filaments and analysed in Imaris software (Bitplane, Zurich, Switzerland). For spine subtype classification custom parameters were used. Classification was entirely automated until the final step where blatant errors in classification were removed.

Time-gated STED imaging was carried out on a Leica TCS SP8 STED 3x microscope running LAS X (Version 2.01.14392) acquisition software using a 100x HC PL APO CS2 oil immersion objective (NA 1.4). Oregon Green 488 (Thermo Fisher Scientific) and Abberoir Star 440SX (Sigma) were excited using the 488nm line (15%) and the 405nm output (20%) from a white light laser (WLL, operating at 70% of its nominal power) respectively. Fluorescence depletion, and therefore super-resolution, was accomplished using a 592nm STED laser (1.5W nominal power, 40% for both Oregon Green 488 and Abberoir Star 440SX). All 2048 × 2048 pixel single plane images were acquired at a scan speed of 400 Hz in bidirectional scan mode. The pixel size of 30.4nm^2^ was optimized for STED imaging. The fluorescence signal was then detected by a Hybrid Detector (HyD, Standard mode) after passing through an Acousto-Optical Beam Splitter (AOBS, detection range 482 - 510nm for Abberoir Star 440SX and 520 - 565nm for Oregon Green 488 when doing two-colour imaging). Time-gated detection was turned on to further improve the resolution in the STED images (0.5 - 6.0ns). The detector gain was adjusted so that no saturation occurred in the images. All 2D STED images were deconvolved using the CMLE (Classic Maximum Likelihood Estimation) algorithm in SVI Huygens Professional (Version 15.10.1p2) to improve the signal-to-noise ratio.

### Electrophysiology in Dissociated Cultures

Whole-cell recordings were performed on transfected cultured hippocampal neurons at 14-17 DIV. Neurons were held at −70 mV. Patch electrodes (4-5 MΩ) were filled with an internal solution containing (in mM): 120 CsCl, 5 QX314 Br, 8 NaCl, 0.2 MgCl_2_, 10 HEPES, 2 EGTA, 2 MgATP and 0.3 Na_3_GTP. The osmolarity and pH were adjusted to 300 mOsm/L and 7.2 respectively. The external artificial cerebro-spinal fluid (ACSF) solution consisted of the following (in mM): 125 NaCl, 25 NaHCO_3_, 2.5 KCl, 2 MgCl_2_, 1.25 NaH_2_PO_4_, 2 CaCl_2_, and 25 glucose saturated with 95% O_2_/5% CO_2_ (pH 7.4, 320 mOsm). This solution was supplemented with CNQX (20 μM), APV (50 μM) and TTX (1μM) to isolate mIPSCs or with bicuculline (20 μM) and TTX (1 μM) for mEPSCs recording. All recordings were performed at room temperatures (22-25 °C). The access resistance, monitored throughout the experiments, was <20 MΩ and results were discarded if it changed by more than 20%. Miniature events and theirs kinetics were analysed using template-based event detection in Clampfit (Molecular Devices, Sunnyvale, CA, USA). Total charge transfer was calculated as described by Peden and colleagues (Peden et al., 2008). For all electrophysiological experiments, the experimenter was blind to the condition/genotype of the sample analysed.

### Acute Hippocampal Slice Electrophysiology

To prepare acute hippocampal slices, male and female mice aged postnatal day 28-34 were used. Immediately after decapitation, the brain was removed and kept in ice-cold dissecting solution. Transverse hippocampal slices (300 μm) were obtained using a vibratome (Leica, VT-1200S). Slices were stored at 35°C for 30 min after slicing and then at 22°C. For the dissection and storage of slices, the solution contained (in mM): 87 NaCl, 25 NaHCO_3_, 10 glucose, 75 sucrose, 2.5 KCl, 1.25 NaH_2_PO_4_, 0.5 CaCl_2_ and 7 MgCl_2_ saturated with 95% O2/5% CO_2_. For patch-clamp experiments, CA1 pyramidal neurons were identified under infrared-differential interference contrast (DIC) imaging with a water-immersion 60X objective (Olympus) and whole-cell recordings were performed as described above for cultured cells.

### Statistics

All data were obtained using cells from at least three independent preparations. Repeats for experiments are given in the figure legends as N numbers and refer to number of cells unless otherwise stated. All statistical analysis was carried out using GraphPad Prism (GraphPad Software, CA, USA) or Microsoft Excel. Data was tested for normal distribution with D’Agostino and Person to determine the use of parametric (unpaired student’s t-test, one-way ANOVA, two-way ANOVA) or non-parametric (Mann-Whitney, Kruskal-Wallis) tests. Appropriate post-hoc tests were carried out in analyses with multiple comparisons and are stated in figure legends. Data are shown as mean ± standard error of the mean (SEM).

## Authors Contributions

Experiments were performed by E.C.D., B.S., J.D. and G.L. Molecular biology was performed by E.C.D and N.F.H. Data were analysed by E.C.D., B.S., J.D., J.T and J.M. and the manuscript was written by E.C.D and J.T.K.

## Acknowledgements

This work was supported by grants to J. T. K. from the UK Medical Research Council (G0802377, MR/N025644/1), BBSRC (BB/I00274X/1) and ERC (282430). E.C.D. was on the MRC LMCB PhD programme. J.D is on the MRC funded 4-year Clinical Neurosciences PhD programme. We thank the UCL Super-resolution Facility (funded by the MRC Next Generation Optical Microscopy Initiative) and MRC LMCB Light Microscopy staff for their contribution.

The authors declare no conflict of interest.

---

**Figure S1:**
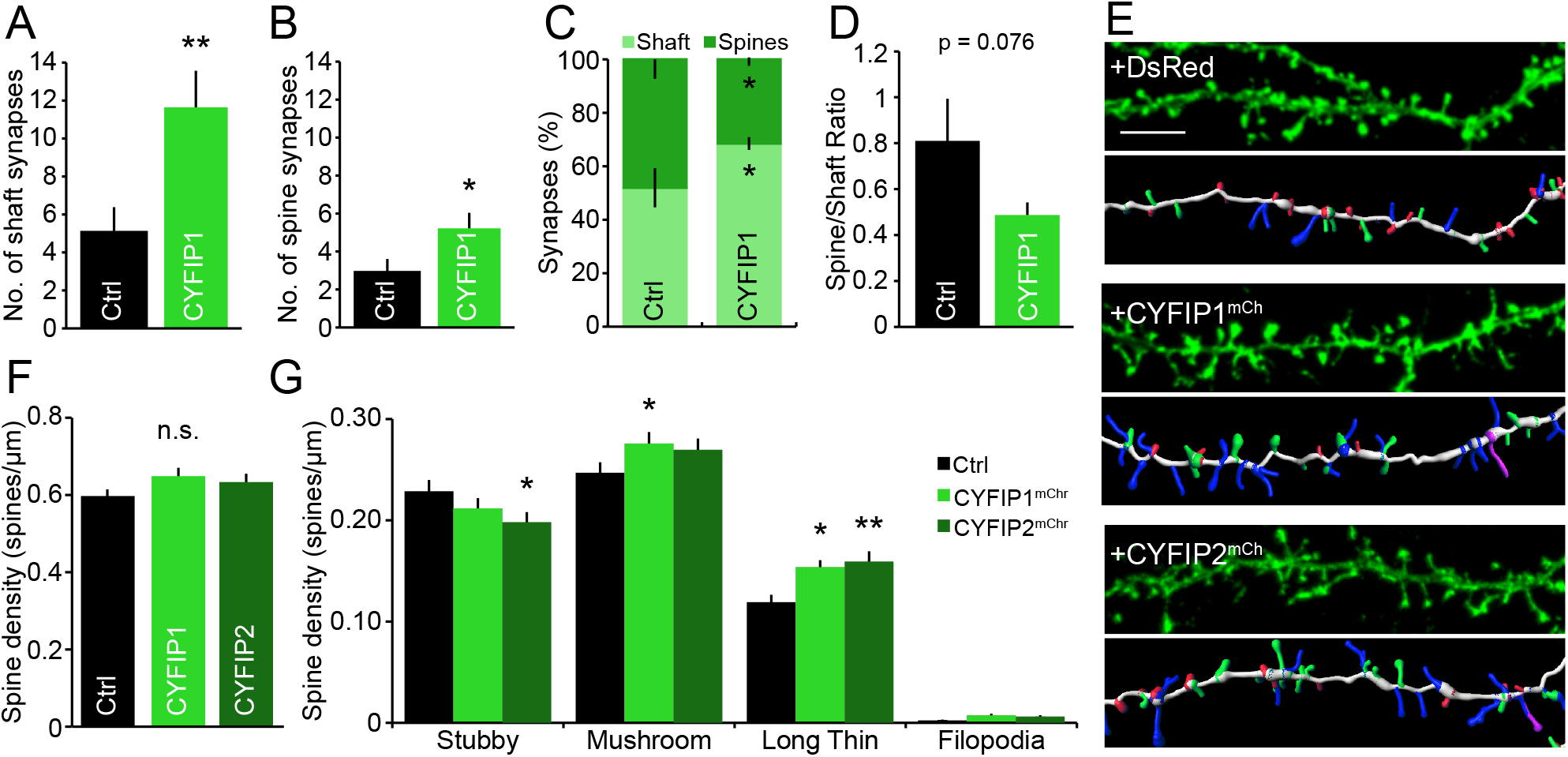
CYFIP1 overexpression redistributes excitatory synapses between dendritic spines and shaft, impacts spine morphology but does not alter spine density. **(A-B)** CYFIP1^GFP^ and GFP control overexpressing neurons were fixed and stained at DIV14 with antibodies to the pre and post excitatory synaptic markers VGLUT and PSD95. Number of synapses, considered as PSD95 and VGLUT positive clusters, were quantified within (A) the dendritic shaft and (B) dendritic spines. CYFIP1^GFP^ overexpression resulted in a significant increase in number of synapses on both the shaft and spines (shaft synapses: from 5.1 ± 1.2 to 11.6 ± 1.9 for CYFIP1, p = 0.0084; spine synapses: from 2.9 ± 0.6 to 5.2 ± 0.8 for CYFIP1, p = 0.039; both n = 15-16 cells from 3 independent preparations; Student’s t-test). **(C)** Graph representing the proportion of excitatory synapses located on dendritic spines compared to the dendritic shaft (shaft synapses: from 52 ± 7.3% to 68.4 ± 2.4% for CYFIP1, n = 14 cells from 3 independent preparations, p = 0.036, Student’s t-test). **(D)** The ratio of spine to shaft excitatory synapses in control compared to CYFIP1^GFP^ overexpressing cells (from 0.8 ± 0.2 to 0.5 ± 0.1, n = 14 cells from 3 independent preparations, p = 0.076 n.s., Student’s t-test). **(E)** Mature hippocampal neurons were transfected for 4 days with actin^GFP^ to label cell morphology and DsRed control, CYFIP1^mCherry^ or CYFIP2^mCherry^, fixed and imaged (upper panels: representative images; lower panels: 3D reconstruction). Colour key for spine 3D reconstruction: green = mushroom, red = stubby, blue = long and thin, pink = filopodia. Scale bar, 5 μm. **(F)** Dendritic spine analysis revealed no change in total spine density (from 0.6 ± 0.02 to 0.65 ± 0.02 for CYFIP1 and 0.63 ± 0.02 for CYFIP2; n=49-66 filaments per condition; 1-way ANOVA, Dunn’s post-hoc multiple comparison, n.s.). **(G)** Quantification of spine subtype density in CYFIP1^mCherry^ and CYFIP2^mCherry^ overexpressing cells (Spines/μm, stubby: from 0.23 ± 0.01 to 0.21 ± 0.01 for CYFIP1 and 0.20 ± 0.01 for CYFIP2; mushroom: from 0.25 ± 0.01 to 0.28 ± 0.01 for CYFIP1 and 0.27 ± 0.01 for CYFIP2; long,thin: from 0.12 ± 0.01 to 0.15 ± 0.01 for CYFIP1 and 0.16 ± 0.01 for CYFIP2; filopodia: from 0.002 ± 0.001 to 0.007 ± 0.002 for CYFIP1 and 0.006 ± 0.001 for CYFIP2; n = 49-66 filaments per condition; 2-way ANOVA, Tukey’s post-hoc multiple comparison). *p < 0.05, **p < 0.01. Bars indicate mean and error bars s.e.m.

**Figure S2:**
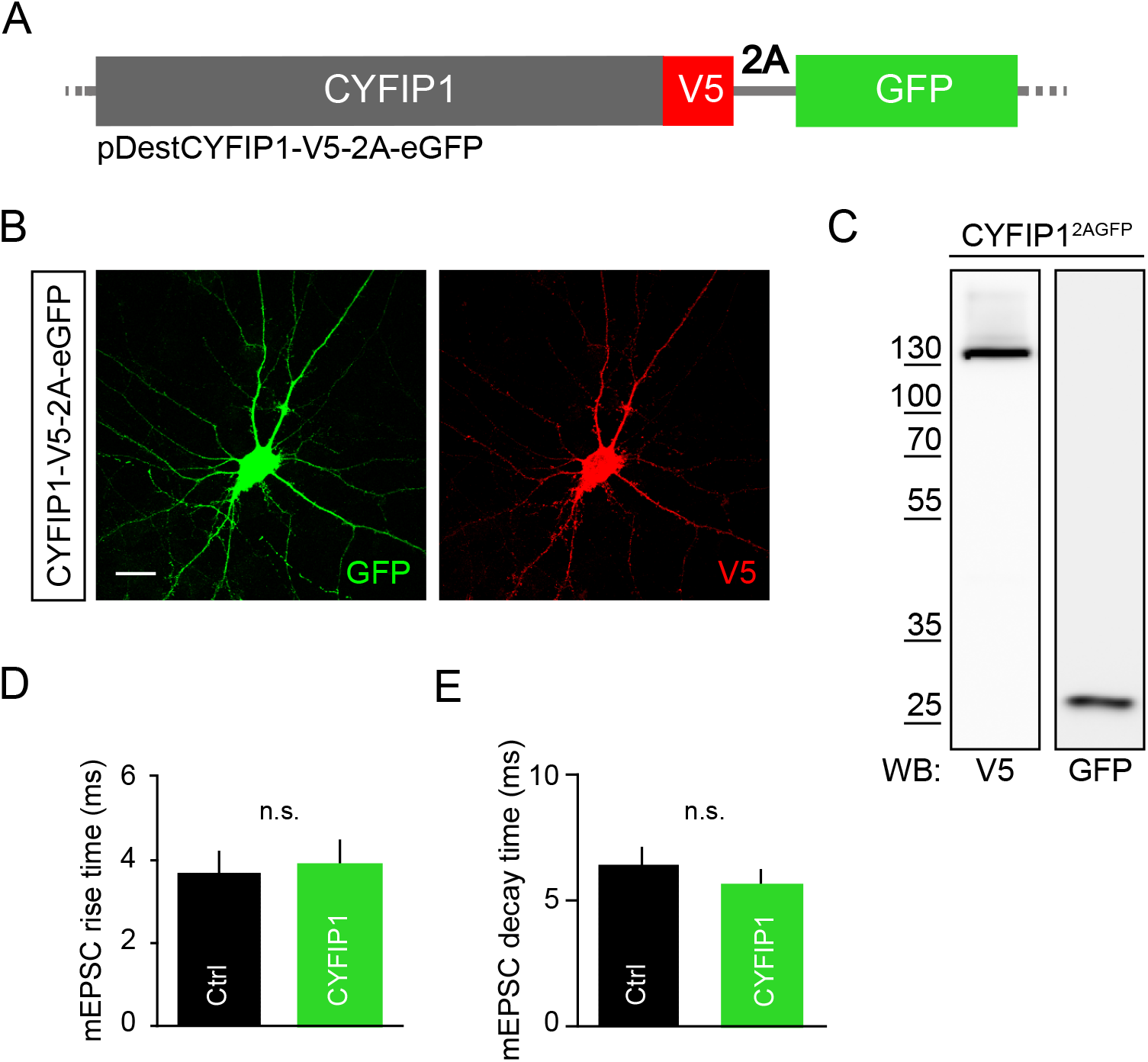
Development and application of the CYFIP1^2AGFP^ construct. To facilitate live cell identification of CYFIP1 overexpressing cells for electrophysiological analysis, CYFIP1 was dually expressed with GFP from a plasmid by the addition of a 2A sequence between the CYFIP1 and GFP cDNA. **(A)** Schematic showing CYFIP1 in the 2A vector. Expression of this vector produces CYFIP1 fused to the small V5 tag and independent expression of cytosolic GFP. Expression of CYFIP1 and GFP is under the same CMV promoter. The 2A sequence in the mRNA causes the transcribing ribosome to skip resulting in the termination of the initial transcribed CYFIP1 sequence and the formation of a new polypeptide at the start of the GFP sequence. **(B)** Hippocampal cells transfected with CYFIP1^2AGFP^. Cells were labelled with antibodies to V5 to confirm CYFIP1 expression and GFP to amplify the cytosolic GFP expression. Scale bar, 20 μm. **(C)** Western blot of COS7 cell lysate sample transfected with CYFIP1^2AGFP^ and probed with antibodies to V5 and GFP. Bands are detected at the expected weight for GFP alone and CYFIP1^V5^ indicating that ribosome skipping and protein expression is occurring correctly. **(D,E)** mEPSCs were recorded from DIV14-16 neurons overexpressing CYFIP1^2AGFP^ or GFP. Calculation of rise (D) and decay (E) time kinetics from these recordings showed no change between GFP and CYFIP1 overexpressing cells (rise time: from 3.7 ± 0.5 ms to 3.9 ± 0.6 ms; n = 12 cells from 3 preparations; p = 0.701 n.s.; Mann-Whitney; decay time: from 6.4 ± 0.7 ms to 5.6 ± 0.6 ms; n = 12 cells from 3 preparations; p = 0.387 n.s.; Student’s t-test).

**Figure S3:**
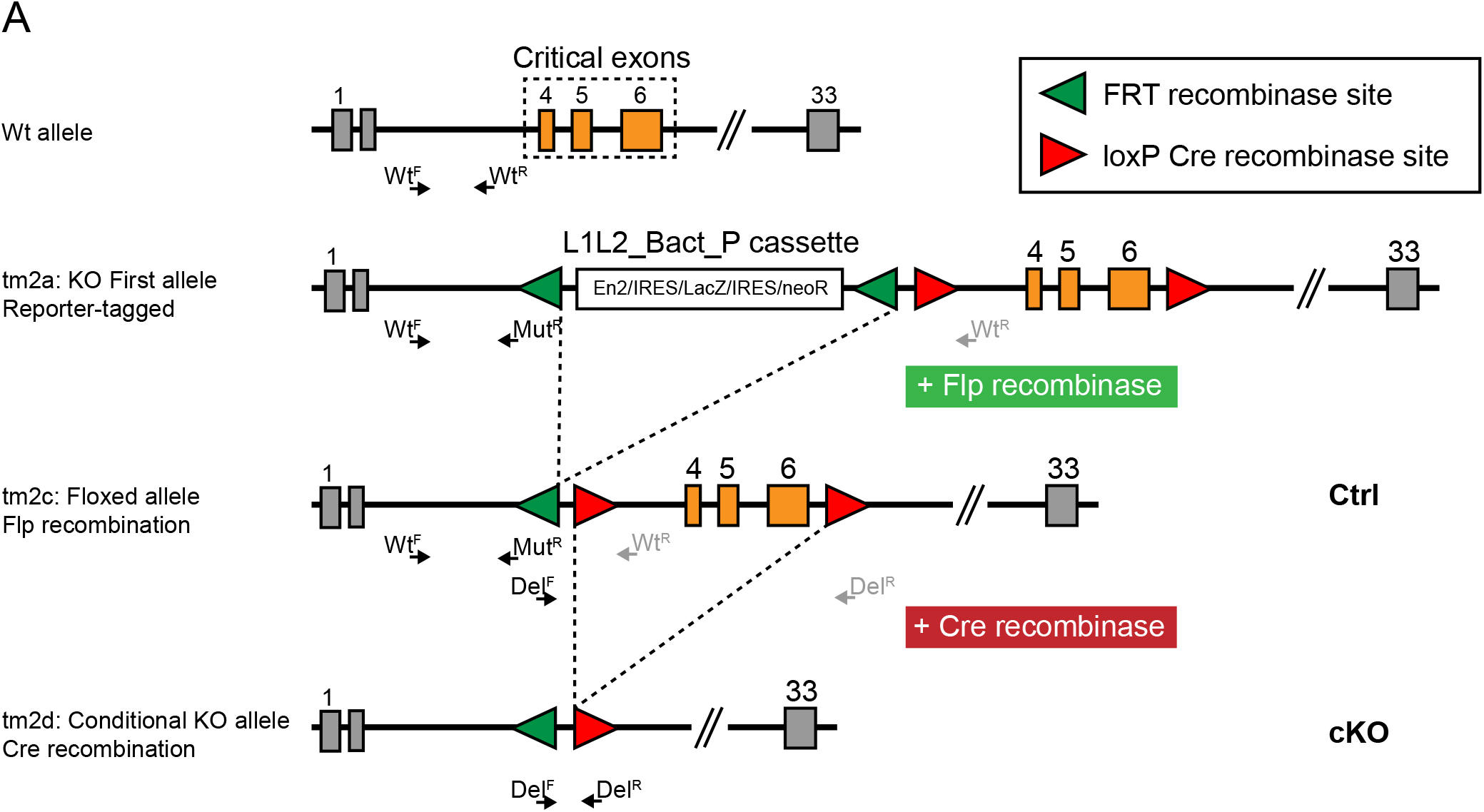
Development and characterisation of the CYFIP1 conditional knockout mouse. **(A)** A schematic of the knockout (KO)-first allele system, demonstrating the generation of the *Cyfip1* floxed allele (tm2c) following Flp recombination of the KO-first allele (tm2a) and then finally the formation of the conditional KO (cKO) allele (tm2d) following Cre recombination of the floxed allele. The KO-first allele contains an IRES:*lacZ* trapping cassette and a promoter-driven *neo* cassette inserted 5’ of critical exons 4 to 6 of *Cyfip1*, disrupting gene function. The cassettes are bound by two frt sites (green triangles) and are extruded in the presence of Flp recombinase, generating the floxed allele. The critical exons are flanked by loxP sites (red triangles), by subjecting the floxed allele to Cre recombination the cKO allele is achieved (Skarnes et al., 2011; White et al., 2013). Primer pair WT^F^ and WT^R^ produce a PCR product of 259 base pairs (bp) from the WT allele, these primers are too distant from each other to produce a product from the KO-first allele and produce a shifted ‘ghost band’ from the floxed allele. Primers WT^F^ and Mut^R^ produced a 182 bp product from the KO-first allele with Mut^R^ annealing at the very 5’ region of the *LacZ* cassette. Primers Del^F^ and Del^R^ produced a 499 bp product from the cKO allele but are too distant from each other to produce a product from the floxed allele.

## Supplemental Experimental Procedures

### Animals

The *Cyfip1* mouse line (MDCK; EPD0555_2_B11; Allele: Cyfip1^tm2a(EUCOMM)Wtsl^) was obtained from the Wellcome Trust Sanger Institute as part of the International Knockout Mouse Consortium (IKMC). Transgenic floxed animals were generated following the Knockout-First strategy on C57BL/6N Taconic USA background (see Supplemental Figure 2) (Skarnes et al., 2011; White et al., 2013). To generate the *Cyfip1* conditional knockout line with ablation of *Cyfip1* expression in the principal cells of the neocortex floxed *Cyfip1* mice were crossed with the Nex-Cre driver line (Goebbels et al., 2006).

Animals were maintained under controlled conditions (temperature 20 ± 2°C; 12 hour light-dark cycle). Food and water were provided *ad libitum*. The genotyping was carried out following Sanger’s recommended procedures, briefly the DNA was extracted from ear biopsies and PCRs were performed with the following primers:

CAS_R1_Term (Mut^R^): TCGTGGTATCGTTATGCGCC
Cyfip1_234230_F (Wt^F^): TGGAAGTAATGGAACCGAACA
Cyfip1_234230_R (Wt^R^): GTAACTACCTATAATGCAGACCTGAAG
Deletion_F (Del^F^): TGGTAGCCCTCTTCTTGTGGA
Deletion_R (Del^R^): CTCCAAGATTCCCCCAAAAC

Control CYFIP1 floxed (CYFIP1^F/F^) and conditional CYFIP1 knock-out animals (CYFIP1^Δ/Δ:Cre^) were generated from CYFIP 1^Δ/Δ^; NEX^Cre^(+/-) x CYFIP 1^F/F^; NEX^Cre^(-/-) crosses. Both male and female mice were used. WT Sprague-Dawley rats were maintained under the same conditions. All procedures for the care and treatment of animals were in accordance with the Animals (Scientific Procedures) Act 1986.

### Constructs

Human CYFIP1- and CYFIP2-GFP/mCherry fusion protein constructs were generated by cloning the coding sequences into pDEST47GFP (Invitrogen) and pDEST-mCherry-N1 (Addgene, 31907) using the Gateway Cloning System (Invitrogen). For CYFIP1-2AGFP the destination vector pDEST-V5:2A:GFP was developed in house. pEGFP-C1 and pCAG-DsRed were purchased from Clontech and Addgene (#24001) respectively. The GFP-actin DNA was a kind gift from J. Hanley (University of Bristol, Bristol, UK).

### Antibodies

Primary antibodies were rabbit anti-CYFIP1 (Upstate, 07-531; ICC, 1:200; WB, 1:1000), mouse anti-GABA_A_R-β2/3 (Neuromab, MAB341; WB, 1:500), guinea pig anti-GABA_A_R-γ2 (Synaptic Systems, 224 004; ICC, 1:500), mouse anti-gephyrin (Synaptic Systems, 147 011; ICC, 1:500; IHC, 1:500; WB, 1:500), rat anti-GFP (Nacalai-Tesque, GF090R; ICC, 1:1000), mouse anti-GFP (Neuromab, 73-131; WB, 1:100), mouse anti-GIT1 (Neuromab, N39/B8; WB, 1:500), rabbit anti-Homer (Synaptic Systems, 160 002; ICC, 1:500; WB, 1:500), rabbit anti-neuroligin 2 (Synaptic Systems, 129 202; WB, 1:1000), mouse anti-neuroligin 3 (Neuromab, N110/29; WB, 1:100), mouse anti-PSD95 (Neuromab, K28/43; ICC, 1:500; WB, 1:1000), rabbit anti-β-PIX (Upstate, 07-1450; WB, 1:2000), mouse anti-β-tubulin (Sigma, M4863-T5293; WB, 1:1000), rabbit anti-vGAT (Synaptic Systems, 131003; ICC, 1:1000), guinea pig anti-vGLUT (Synaptic Systems, 135304; ICC, 1:1000), mouse anti-V5 (Invitrogen, R960-25, ICC, 1:1000; WB, 1:1000). Secondary fluorescent antibodies were conjugated to Alexa Fluor 488, 568, 647 (1:1000, Molecular Probes). Anti-rabbit and anti-mouse HRP conjugated secondaries were from Jackson ImmunoResearch (WB, 1;1000).

### Cell Culture and Transfections

Hippocampal cultures were obtained from either E18 Sprague Dawley WT rat or E16 mouse embryos (produced via CYFIP1^Δ/Δ^; NEX^Cre^(+/-) x CYFIP1^F/F^; NEX^Cre^(-/-) crosses) of either sex as previously described (López-Doménech et al., 2016; Vaccaro et al., 2017). Neurons were transfected using Lipofectamine-2000 (Invitrogen). COS-7 cells were maintained in DMEM supplemented with fetal calf serum and antibiotics and transfected using the Nucleofector^®^ device (Amaxa) following the manufacturer’s protocol.

### Proximity Ligation Assay

Proximity ligation assays (Duolink) were carried out using anti-CYFIP1 and anti-gephyrin antibodies or anti-gephyrin alone for control proximity ligation assays. Neurons were fixed and blocked as for immunofluorescence and incubated with primary antibodies. Following primary antibody incubation, cells were washed in PBS before incubation with secondary antibodies conjugated with oligonucleotides.

Ligation and amplification reactions were conducted at 37 °C, as described in the Duolink manual, before mounting and visualization using confocal microscopy (Norkett et al., 2015). For PLA analysis, confocal image stacks with a X0.5 zoom and voxel dimensions 0.39 μm x 0.39 μm x 0.57 μm were acquired. Analysis was carried out on maximum projection images using Metamorph (Molecular Devices, Sunnyvale, CA, USA). A user-defined threshold was applied to each image which best detected PLA puncta and kept constant within an experiment. Puncta were then counted per field of view.

### Preparation of Brain Lysates

Adult WT and conditional KO male or female whole brains or cortical regions were sonicated in lysis buffer (50 mM HEPES pH 7.5, 0.5 % Triton-X-100, 150 mM NaCl, 1 mM EDTA, 1 mM PMSF in the presence of antipain, pepstatin and leupeptin) then left to rotate at 4 °C for 1 hour. Membranes were pelleted by centrifugation at 14000 g for 15 minutes at 4 °C. Protein content of the supernatant was assayed by BioRad protein assay. Samples were then suspended in 3X protein sample buffer and analyzed by SDS-PAGE and western blotting. Briefly, protein samples were separated by SDS-PAGE on 10 % Tris-Glycine gels and blotted onto nitrocellulose membranes (GE Healthcare Bio-Sciences). Membranes were blocked for 1 hour in milk (PBS, 0.05 % Tween, 4 % milk), incubated in primary antibodies diluted with milk overnight at 4 °C before incubation in an appropriate HRP-conjugated secondary antibody for 1 hour at room temperature. The blots were developed with an ECL-Plus detection reagent (GE Healthcare Bio-Sciences). Densitometric analysis was performed in ImageJ (NIH).

### Golgi Staining

Dendritic and spine morphology in P30 mice was analysed using the FD Rapid Golgi Stain kit (FD NeuroTechnologies, Baltimore, MD, USA) and Neurolucida (MBF Bioscience, Williston, VT, USA). Golgi-impregnated brains were sliced at 100μm using a vibratome (Leica Microsystems, Heerbrugg, Switzerland). Well-isolated hippocampal CA1 neurons were imaged at 20X using the Neurolucida software system and an upright light microscope with a motorized stage (MBF Bioscience). The entire dendritic tree (apical and basal) was traced and reconstructed. 3-dimensional Sholl analysis of reconstructions was performed using a custom MATLAB script. For spine analysis, 50μm z-stacks of 2μm step size were imaged at 40X using a ZEISS Axio Scan system and sections of basal dendrite were randomly selected for analysis. Spine length and head width were manually traced in ImageJ and the data analysed using a custom EXCEL macro.

## References

Abekhoukh, S., Sahin, H.B., Grossi, M., Zongaro, S., Maurin, T., Madrigal, I., Kazue-Sugioka, D., Raas-Rothschild, A., Doulazmi, M., Carrera, P., Stachon, A., Scherer, S., Drula Do Nascimento, M.R., Trembleau, A., Arroyo, I., Szatmari, P., Smith, I.M., Milà, M., Smith, A.C., et al. (2017). New insights into the regulatory function of CYFIP1 in the context of WAVE- and FMRP-containing complexes. Dis. Model. Mech. 10, 463–474.

Anazi, S., Maddirevula, S., Salpietro, V., Asi, Y.T., Alsahli, S., Alhashem, A., Shamseldin, H.E., AlZahrani, F., Patel, N., Ibrahim, N., Abdulwahab, F.M., Hashem, M., Alhashmi, N., Al Murshedi, F., Al Kindy, A., Alshaer, A., Rumayyan, A., Al Tala, S., Kurdi, W., et al. (2017). Expanding the genetic heterogeneity of intellectual disability. Hum. Genet. 136, 1419–1429.

Anitei, Stange, Parshina, Baust, Schenck, Raposo, Kirchhausen, and Hoflack (2010). Protein complexes containing CYFIP/Sra/PIR121 coordinate Arf1 and Rac1 signalling during clathrin-AP-1-coated carrier biogenesis at the TGN. Nat. Publ. Gr. 12, 330–340.

Bannai, H., Lévi, S., Schweizer, C., Inoue, T., Launey, T., Racine, V., Sibarita, J.-B., Mikoshiba, K., and Triller, A. (2009). Activity-dependent tuning of inhibitory neurotransmission based on GABAAR diffusion dynamics. Neuron 62, 670–682.

Bateup, H.S., Johnson, C.A., Denefrio, C.L., Saulnier, J.L., Kornacker, K., and Sabatini, B.L. (2013). Excitatory/inhibitory synaptic imbalance leads to hippocampal hyperexcitability in mouse models of tuberous sclerosis. Neuron 78, 510–522.

Bemben, M.A., Shipman, S.L., Nicoll, R.A., and Roche, K.W. (2015a). The cellular and molecular landscape of neuroligins. Trends Neurosci. 38, 496–505.

Bemben, M.A., Nguyen, Q.-A., Wang, T., Li, Y., Nicoll, R.A., and Roche, K.W. (2015b). Autism-associated mutation inhibits protein kinase C-mediated neuroligin-4X enhancement of excitatory synapses. Proc. Natl. Acad. Sci. 112, 2551–2556.

Blundell, J., Tabuchi, K., Bolliger, M.F., Blaiss, C.A., Brose, N., Liu, X., Südhof, T.C., and Powell, C.M. (2009). Increased anxiety-like behavior in mice lacking the inhibitory synapse cell adhesion molecule neuroligin 2. Genes. Brain. Behav. 8, 114–126.

Bourgeron T. (2015). From the genetic architecture to synaptic plasticity in autism spectrum disorder. Nat. Rev. Neurosci. 16, 551–563.

Bozdagi, O., Sakurai, T., Dorr, N., Pilorge, M., Takahashi, N., and Buxbaum, J.D. (2012). Haploinsufficiency of cyfip1 produces fragile x-like phenotypes in mice. PLoS One 7, e42422.

Budreck, E.C., and Scheiffele, P. (2007). Neuroligin-3 is a neuronal adhesion protein at GABAergic and glutamatergic synapses. Eur. J. Neurosci. 26, 1738–1748.

Chanda, S., Hale, W.D., Zhang, B., Wernig, M., and Südhof, T.C. (2017). Unique versus Redundant Functions of Neuroligin Genes in Shaping Excitatory and Inhibitory Synapse Properties. J. Neurosci. 37, 6816–6836.

Chao, H.-T., Chen, H., Samaco, R.C., Xue, M., Chahrour, M., Yoo, J., Neul, J.L., Gong, S., Lu, H.-C., Heintz, N., Ekker, M., Rubenstein, J.L.R., Noebels, J.L., Rosenmund, C., and Zoghbi, H.Y. (2010). Dysfunction in GABA signalling mediates autism-like stereotypies and Rett syndrome phenotypes. Nature 468, 263–269.

Chen, B., Brinkmann, K., Chen, Z., Pak, C.W., Liao, Y., Shi, S., Henry, L., Grishin N. V, Bogdan, S., and Rosen, M.K. (2014). The WAVE regulatory complex links diverse receptors to the actin cytoskeleton. Cell 156, 195–207.

Chen, Z., Borek, D., Padrick, S.B., Gomez, T.S., Metlagel, Z., Ismail, A.M., Umetani, J., Billadeau, D.D., Otwinowski, Z., and Rosen, M.K. (2010). Structure and control of the actin regulatory WAVE complex. Nature 468, 533–538.

Chia, P.H., Chen, B., Li, P., Rosen, M.K., and Shen, K. (2014). Local F-actin network links synapse formation and axon branching. Cell 156, 208–220.

Chung, L., Wang, X., Zhu, L., Towers, A., Kim, I.H., and Jiang, Y.-H. (2015). Parental origin impairment of synaptic functions and behaviors in cytoplasmic FMRP interacting protein 1 (Cyfip1) deficiency mice. Brain Res. 1629, 340–350.

Crestani, F., Lorez, M., Baer, K., Essrich, C., Benke, D., Laurent, J.P., Belzung, C., Fritschy, J.M., Lüscher, B., and Mohler, H. (1999). Decreased GABAA-receptor clustering results in enhanced anxiety and a bias for threat cues. Nat. Neurosci. 2, 833–839.

Darnell, J.C., Van Driesche, S.J., Zhang, C., Hung, K.Y.S., Mele, A., Fraser, C.E., Stone, E.F., Chen, C., Fak, J.J., Chi, S.W., Licatalosi, D.D., Richter, J.D., and Darnell, R.B. (2011). FMRP stalls ribosomal translocation on mRNAs linked to synaptic function and autism. Cell 146, 247–261.

Davenport, E.C., Pendolino, V., Kontou, G., McGee, T.P., Sheehan, D.F., López-Doménech, G., Farrant, M., and Kittler, J.T. (2017). An Essential Role for the Tetraspanin LHFPL4 in the Cell-Type-Specific Targeting and Clustering of Synaptic GABA A Receptors. Cell Rep. 21, 7083.

Doornbos M., Sikkema-Raddatz B., Ruijvenkamp C. a L., Dijkhuizen, T., Bijlsma, E.K., Gijsbers, A.C.J., Hilhorst-Hofstee, Y., Hordijk, R., Verbruggen, K.T., Kerstjens-Frederikse, W.S. (Mieke), van Essen, T., Kok, K., van Silfhout, A.T., Breuning, M., and van Ravenswaaij-Arts, C.M. a (2009). Nine patients with a microdeletion 15q11.2 between breakpoints 1 and 2 of the Prader-Willi critical region, possibly associated with behavioural disturbances. Eur. J. Med. Genet. 52, 108–115.

Fekete, C.D., Chiou, T.-T., Miralles, C.P., Harris, R.S., Fiondella, C.G., Loturco, J.J., and De Blas, A.L. (2015). In vivo clonal overexpression of neuroligin 3 and neuroligin 2 in neurons of the rat cerebral cortex: Differential effects on GABAergic synapses and neuronal migration. J. Comp. Neurol. 523, 1359–1378.

Föcking, M., Lopez, L.M., English, J.A., Dicker, P., Wolff, A., Brindley, E., Wynne, K., Cagney, G., and Cotter, D.R. (2014). Proteomic and genomic evidence implicates the postsynaptic density in schizophrenia. Mol. Psychiatry.

Foss-Feig, J.H., Adkinson, B.D., Ji, J.L., Yang, G., Srihari, V.H., McPartland, J.C., Krystal, J.H., Murray, J.D., and Anticevic, A. (2017). Searching for Cross-Diagnostic Convergence: Neural Mechanisms Governing Excitation and Inhibition Balance in Schizophrenia and Autism Spectrum Disorders. Biol. Psychiatry 81, 848–861.

Fromer, M., Pocklington, A.J., Kavanagh, D.H., Williams, H.J., Dwyer, S., Gormley, P., Georgieva, L., Rees, E., Palta, P., Ruderfer, D.M., Carrera, N., Humphreys, I., Johnson, J.S., Roussos, P., Barker, D.D., Banks, E., Milanova, V., Grant, S.G., Hannon, E., et al. (2014). De novo mutations in schizophrenia implicate synaptic networks. Nature 506, 179–184.

Gao, R., and Penzes, P. (2015). Common mechanisms of excitatory and inhibitory imbalance in schizophrenia and autism spectrum disorders. Curr. Mol. Med. 15, 146–167.

Gkogkas, C.G., Khoutorsky, A., Ran, I., Rampakakis, E., Nevarko, T., Weatherill, D.B., Vasuta, C., Yee, S., Truitt, M., Dallaire, P., Major, F., Lasko, P., Ruggero, D., Nader, K., Lacaille, J.-C., and Sonenberg, N. (2013). Autism-related deficits via dysregulated eIF4E-dependent translational control. Nature 493, 371–377.

Goebbels, S., Bormuth, I., Bode, U., Hermanson, O., Schwab, M.H., and Nave, K.-A. (2006). Genetic targeting of principal neurons in neocortex and hippocampus of NEX-Cre mice. Genesis 44, 611–621.

Han, K., Chen, H., Gennarino, V.A., Richman, R., Lu, H.-C., and Zoghbi, H.Y. (2015). Fragile X-like behaviors and abnormal cortical dendritic spines in Cytoplasmic FMR1-interacting protein 2-mutant mice. Hum. Mol. Genet. 24, 1813–1823.

Hoeffer, C.A., Sanchez, E., Hagerman, R.J., Mu, Y., Nguyen D. V, Wong, H., Whelan, A.M., Zukin, R.S., Klann, E., and Tassone, F. (2012). Altered mTOR signaling and enhanced CYFIP2 expression levels in subjects with fragile X syndrome. Genes. Brain. Behav. 11, 332–341.

Horn, M.E., and Nicoll, R.A. (2018). Somatostatin and parvalbumin inhibitory synapses onto hippocampal pyramidal neurons are regulated by distinct mechanisms. Proc. Natl. Acad. Sci. U. S. A. 115, 589–594.

Hsiao, K., Harony-Nicolas, H., Buxbaum, J.D., Bozdagi-Gunal, O., and Benson, D.L. (2016). Cyfip1 Regulates Presynaptic Activity during Development. J. Neurosci. 36, 1564–1576.

Huang Y. (2015). Up-Regulated Cytoplasmic FMRP-interacting protein 1 in Intractable Temporal Lobe Epilepsy Patients and a Rat Model. Int. J. Neurosci. 1–28.

Iossifov, I., O’Roak, B.J., Sanders, S.J., Ronemus, M., Krumm, N., Levy, D., Stessman, H.A., Witherspoon, K.T., Vives, L., Patterson, K.E., Smith, J.D., Paeper, B., Nickerson, D.A., Dea, J., Dong, S., Gonzalez, L.E., Mandell, J.D., Mane, S.M., Murtha, M.T., et al. (2014). The contribution of de novo coding mutations to autism spectrum disorder. Nature 515, 216–221.

Jamain, S., Quach, H., Betancur, C., Råstam, M., Colineaux, C., Gillberg, I.C., Soderstrom, H., Giros, B., Leboyer, M., Gillberg, C., Bourgeron, T., and Paris Autism Research International Sibpair Study (2003). Mutations of the X-linked genes encoding neuroligins NLGN3 and NLGN4 are associated with autism. Nat. Genet. 34, 27–29.

Kim, J.H., Lee, S.-R., Li, L.-H., Park, H.-J., Park, J.-H., Lee, K.Y., Kim, M.-K., Shin, B.A., and Choi, S.-Y. (2011). High cleavage efficiency of a 2A peptide derived from porcine teschovirus-1 in human cell lines, zebrafish and mice. PLoS One 6, e18556.

Kirkpatrick, S.L., Goldberg, L.R., Yazdani, N., Babbs, R.K., Wu, J., Reed, E.R., Jenkins, D.F., Bolgioni, A.F., Landaverde, K.I., Luttik, K.P., Mitchell, K.S., Kumar, V., Johnson, W.E., Mulligan, M.K., Cottone, P., and Bryant, C.D. (2017). Cytoplasmic FMR1-Interacting Protein 2 Is a Major Genetic Factor Underlying Binge Eating. Biol. Psychiatry 81, 757–769.

Kobayashi, K., Kuroda, S., Fukata, M., Nakamura, T., Nagase, T., Nomura, N., Matsuura, Y., Yoshida-Kubomura, N., Iwamatsu, a, and Kaibuchi, K. (1998). p140Sra-1 (specifically Rac1-associated protein) is a novel specific target for Rac1 small GTPase. J. Biol. Chem. 273, 291–295.

de Kovel, C.G.F., Trucks, H., Helbig, I., Mefford, H.C., Baker, C., Leu, C., Kluck, C., Muhle, H., von Spiczak, S., Ostertag, P., Obermeier, T., Kleefuss-Lie, A.A., Hallmann, K., Steffens, M., Gaus, V., Klein, K.M., Hamer, H.M., Rosenow, F., Brilstra, E.H., et al. (2010). Recurrent microdeletions at 15q11.2 and 16p13.11 predispose to idiopathic generalized epilepsies. Brain 133, 23–32.

Kumar, V., Kim, K., Joseph, C., Kourrich, S., Yoo, S.-H., Huang, H.C., Vitaterna, M.H., Pardo-Manuel de Villena, F., Churchill, G., Bonci, A., and Takahashi, J.S. (2013). C57BL/6N Mutation in Cytoplasmic FMRP interacting protein 2 Regulates Cocaine Response. Science (80-.). 342, 1508–1512.

Leblond, C.S., Heinrich, J., Delorme, R., Proepper, C., Betancur, C., Huguet, G., Konyukh, M., Chaste, P., Ey, E., Rastam, M., Anckarsäter, H., Nygren, G., Gillberg, I.C., Melke, J., Toro, R., Regnault, B., Fauchereau, F., Mercati, O., Lemière, N., et al. (2012). Genetic and functional analyses of SHANK2 mutations suggest a multiple hit model of autism spectrum disorders. PLoS Genet. 8, e1002521.

Luscher, B., Fuchs, T., and Kilpatrick, C. (2011). GABAA receptor trafficking-mediated plasticity of inhibitory synapses. Neuron 70, 385–409.

Marshall, C.R., Howrigan, D.P., Merico, D., Thiruvahindrapuram, B., Wu, W., Greer, D.S., Antaki, D., Shetty, A., Holmans, P.A., Pinto, D., Gujral, M., Brandler, W.M., Malhotra, D., Wang, Z., Fajarado, K.V.F., Maile, M.S., Ripke, S., Agartz, I., Albus, M., et al. (2017). Contribution of copy number variants to schizophrenia from a genome-wide study of 41,321 subjects. Nat. Genet. 49, 27–35.

Muir, J., Arancibia-Carcamo, I.L., MacAskill, A.F., Smith, K.R., Griffin, L.D., and Kittler, J.T. (2010). NMDA receptors regulate GABAA receptor lateral mobility and clustering at inhibitory synapses through serine 327 on the γ2 subunit. Proc. Natl. Acad. Sci. U. S. A. 107, 16679–16684.

Nakashima, M., Kato, M., Aoto, K., Shiina, M., Belal, H., Mukaida, S., Kumada, S., Sato, A., Zerem, A., Lerman-Sagie, T., Lev, D., Leong, H.Y., Tsurusaki, Y., Mizuguchi, T., Miyatake, S., Miyake, N., Ogata, K., Saitsu, H., and Matsumoto, N. (2018). De Novo Hotspot Variants in CYFIP2 Cause Early-Onset Epileptic Encephalopathy. Ann. Neurol.

Napoli, I., Mercaldo, V., Boyl, P.P., Eleuteri, B., Zalfa, F., De Rubeis, S., Di Marino, D., Mohr, E., Massimi, M., Falconi, M., Witke, W., Costa-Mattioli, M., Sonenberg, N., Achsel, T., and Bagni, C. (2008). The fragile X syndrome protein represses activity-dependent translation through CYFIP1, a new 4E-BP. Cell 134, 1042–1054.

Nebel, R.A., Zhao, D., Pedrosa, E., Kirschen, J., Lachman, H.M., Zheng, D., and Abrahams, B. S. (2016). Reduced CYFIP1 in Human Neural Progenitors Results in Dysregulation of Schizophrenia and Epilepsy Gene Networks. PLoS One 11, e0148039.

Nelson, S.B., and Valakh, V. (2015). Excitatory/Inhibitory Balance and Circuit Homeostasis in Autism Spectrum Disorders. Neuron 87, 684–698.

Nishimura, Y., Martin, C.L., Vazquez-Lopez, A., Spence, S.J., Alvarez-Retuerto, A.I., Sigman, M., Steindler, C., Pellegrini, S., Schanen, N.C., Warren, S.T., and Geschwind, D.H. (2007). Genome-wide expression profiling of lymphoblastoid cell lines distinguishes different forms of autism and reveals shared pathways. Hum. Mol. Genet. 16, 1682–1698.

Norkett, R., Modi, S., Birsa, N., Atkin, T.A., Ivankovic, D., Pathania, M., Trossbach S. V, Korth, C., Hirst, W.D., and Kittler, J.T. (2015). DISC1-dependent Regulation of Mitochondrial Dynamics Controls the Morphogenesis of Complex Neuronal Dendrites. J. Biol. Chem. 291, 613–629.

Oguro-Ando, A, Rosensweig, C., Herman, E., Nishimura, Y., Werling, D., Bill, B.R., Berg, J.M., Gao, F., Coppola, G., Abrahams, B.S., and Geschwind, D.H. (2014). Increased CYFIP1 dosage alters cellular and dendritic morphology and dysregulates mTOR. Mol. Psychiatry.

Paluszkiewicz, S.M., Martin, B.S., and Huntsman, M.M. (2011). Fragile X syndrome: the GABAergic system and circuit dysfunction. Dev. Neurosci. 33, 349–364.

Pathania, M., Davenport, E.C., Muir, J., Sheehan, D.F., López-Doménech, G., and Kittler, J.T. (2014). The autism and schizophrenia associated gene CYFIP1 is critical for the maintenance of dendritic complexity and the stabilization of mature spines. Transl. Psychiatry 4, e374.

Peden, D.R., Petitjean, C.M., Herd, M.B., Durakoglugil, M.S., Rosahl, T.W., Wafford, K., Homanics, G.E., Belelli, D., Fritschy, J.-M., and Lambert, J.J. (2008). Developmental maturation of synaptic and extrasynaptic GABAA receptors in mouse thalamic ventrobasal neurones. J. Physiol. 586, 965–987.

Pettem, K.L., Yokomaku, D., Luo, L., Linhoff, M.W., Prasad, T., Connor, S.A., Siddiqui, T.J., Kawabe, H., Chen, F., Zhang, L., Rudenko, G., Wang, Y.T., Brose, N., and Craig, A.M. (2013). The specific α-neurexin interactor calsyntenin-3 promotes excitatory and inhibitory synapse development. Neuron 80, 113–128.

Phillips, M., and Pozzo-Miller, L. (2015). Dendritic spine dysgenesis in autism related disorders. Neurosci. Lett. 601, 30–40.

Picinelli, C., Lintas, C., Piras, I.S., Gabriele, S., Sacco, R., Brogna, C., and Persico, A.M. (2016). Recurrent 15q11.2 BP1-BP2 microdeletions and microduplications in the etiology of neurodevelopmental disorders. Am. J. Med. Genet. Part B Neuropsychiatr. Genet. 171, 1088–1098.

Pinto, D., Delaby, E., Merico, D., Barbosa, M., Merikangas, A., Klei, L., Thiruvahindrapuram, B., Xu, X., Ziman, R., Wang, Z., Vorstman, J.A.S., Thompson, A., Regan, R., Pilorge, M., Pellecchia, G., Pagnamenta, A.T., Oliveira, B., Marshall, C.R., Magalhaes, T.R., et al. (2014). Convergence of Genes and Cellular Pathways Dysregulated in Autism Spectrum Disorders. Am. J. Hum. Genet. 94, 677.

Poulopoulos, A., Aramuni, G., Meyer, G., Soykan, T., Hoon, M., Papadopoulos, T., Zhang, M., Paarmann, I., Fuchs, C., Harvey, K., Jedlicka, P., Schwarzacher, S.W., Betz, H., Harvey, R.J., Brose, N., Zhang, W., and Varoqueaux, F. (2009). Neuroligin 2 drives postsynaptic assembly at perisomatic inhibitory synapses through gephyrin and collybistin. Neuron 63, 628–642.

Radyushkin, K., Hammerschmidt, K., Boretius, S., Varoqueaux, F., El-Kordi, A., Ronnenberg, A., Winter, D., Frahm, J., Fischer, J., Brose, N., and Ehrenreich, H. (2009). Neuroligin-3-deficient mice: model of a monogenic heritable form of autism with an olfactory deficit. Genes. Brain. Behav. 8, 416–425.

Rees, E., Walters, J.T.R., Georgieva, L., Isles, A.R., Chambert, K.D., Richards, A.L., Mahoney-Davies, G., Legge, S.E., Moran, J.L., McCarroll, S.A., O’Donovan, M.C., Owen, M.J., and Kirov, G. (2014). Analysis of copy number variations at 15 schizophrenia-associated loci. Br. J. Psychiatry 204, 108–114.

De Rubeis, S., Pasciuto, E., Li, K.W., Fernández, E., Di Marino, D., Buzzi, A., Ostroff, L.E., Klann, E., Zwartkruis, F.J.T., Komiyama, N.H., Grant, S.G.N., Poujol, C., Choquet, D., Achsel, T., Posthuma, D., Smit, A.B., and Bagni, C. (2013). CYFIP1 coordinates mRNA translation and cytoskeleton remodeling to ensure proper dendritic spine formation. Neuron 79, 1169–1182.

De Rubeis, S., He, X., Goldberg, A.P., Poultney, C.S., Samocha, K., Ercument Cicek, A., Kou, Y., Liu, L., Fromer, M., Walker, S., Singh, T., Klei, L., Kosmicki, J., Fu, S.-C., Aleksic, B., Biscaldi, M., Bolton, P.F., Brownfeld, J.M., Cai, J., et al. (2014). Synaptic, transcriptional and chromatin genes disrupted in autism. Nature 515, 209–215.

Rudolph, U., and Möhler, H. (2014). GABAA receptor subtypes: Therapeutic potential in Down syndrome, affective disorders, schizophrenia, and autism. Annu. Rev. Pharmacol. Toxicol. 54, 483–507.

Skarnes, W.C., Rosen, B., West, A.P., Koutsourakis, M., Bushell, W., Iyer, V., Mujica, A.O., Thomas, M., Harrow, J., Cox, T., Jackson, D., Severin, J., Biggs, P., Fu, J., Nefedov, M., de Jong, P.J., Stewart, A.F., and Bradley, A. (2011). A conditional knockout resource for the genome-wide study of mouse gene function. Nature 474, 337–342.

Smith, K.R., Davenport, E.C., Wei, J., Li, X., Pathania, M., Vaccaro, V., Yan, Z., and Kittler, J.T. (2014). GIT 1 and βPIX are essential for GABAA receptor synaptic stability and inhibitory neurotransmission. Cell Rep. 9, 298–310.

Smith, K.R., Jones, K.A., Kopeikina, K.J., Burette, A.C., Copits, B.A., Yoon, S., Forrest, M.P., Fawcett-Patel, J.M., Hanley, J.G., Weinberg, R.J., Swanson, G.T., and Penzes, P. (2017). Cadherin-10 Maintains Excitatory/Inhibitory Ratio through Interactions with Synaptic Proteins. J. Neurosci. 37, 11127–11139.

Stefansson, H., Rujescu, D., Cichon, S., Pietiläinen, O.P.H., Ingason, A., Steinberg, S., Fossdal, R., Sigurdsson, E., Sigmundsson, T., Buizer-Voskamp, J.E., Hansen, T., Jakobsen, K.D., Muglia, P., Francks, C., Matthews, P.M., Gylfason, A., Halldorsson B. V, Gudbjartsson, D., Thorgeirsson, T.E., et al. (2008). Large recurrent microdeletions associated with schizophrenia. Nature 455, 232–236.

Tabuchi, K., Blundell, J., Etherton, M.R., Hammer, R.E., Liu, X., Powell, C.M., and Südhof, T.C. (2007). A neuroligin-3 mutation implicated in autism increases inhibitory synaptic transmission in mice. Science 318, 71–76.

Tam, G.W.C., van de Lagemaat, L.N., Redon, R., Strathdee, K.E., Croning, M.D.R., Malloy, M.P., Muir, W.J., Pickard, B.S., Deary, I.J., Blackwood, D.H.R., Carter, N.P., and Grant, S.G.N. (2010). Confirmed rare copy number variants implicate novel genes in schizophrenia. Biochem. Soc. Trans. 38, 445–451.

Tiwari, S.S., Mizuno, K., Ghosh, A., Aziz, W., Troakes, C., Daoud, J., Golash, V., Noble, W., Hortobágyi, T., and Giese, K.P. (2016). Alzheimer-related decrease in CYFIP2 links amyloid production to tau hyperphosphorylation and memory loss. Brain 139, 2751–2765.

Toma, C., Torrico, B., Hervás, A., Valdés-Mas, R., Tristán-Noguero, A., Padillo, V., Maristany, M., Salgado, M., Arenas, C., Puente, X.S., Bayés, M., and Cormand, B. (2014). Exome sequencing in multiplex autism families suggests a major role for heterozygous truncating mutations. Mol. Psychiatry 19, 784–790.

Tora, D., Gomez, A.M., Michaud, J.-F., Yam, P.T., Charron, F., and Scheiffele, P. (2017). Cellular Functions of the Autism Risk Factor PTCHD1 in Mice. J. Neurosci. 37, 11993–12005.

Twelvetrees, A.E., Yuen, E.Y., Arancibia-Carcamo, I.L., MacAskill, A.F., Rostaing, P., Lumb, M.J., Humbert, S., Triller, A., Saudou, F., Yan, Z., and Kittler, J.T. (2010). Delivery of GABAARs to synapses is mediated by HAP1-KIF5 and disrupted by mutant huntingtin. Neuron 65, 53–65.

Tyagarajan, S.K., and Fritschy, J.-M. (2014). Gephyrin: a master regulator of neuronal function? Nat. Rev. Neurosci. 15, 141–156.

Uezu, A., Kanak, D.J., Bradshaw, T.W.A., Soderblom, E.J., Catavero, C.M., Burette, A.C., Weinberg, R.J., and Soderling, S.H. (2016). Identification of an elaborate complex mediating postsynaptic inhibition. Science (80-.). 353, 1123–1129.

Vanlerberghe, C., Petit, F., Malan, V., Vincent-Delorme, C., Bouquillon, S., Boute, O., Holder-Espinasse, M., Delobel, B., Duban, B., Vallee, L., Cuisset, J.-M., Lemaitre, M.-P., Vantyghem, M.-C., Pigeyre, M., Lanco-Dosen, S., Plessis, G., Gerard, M., Decamp, M., Mathieu, M., et al. (2015). 15q11.2 microdeletion (BP1-BP2) and developmental delay, behaviour issues, epilepsy and congenital heart disease: A series of 52 patients. Eur. J. Med. Genet. 58, 140–147.

Varoqueaux, F., Jamain, S., and Brose, N. (2004). Neuroligin 2 is exclusively localized to inhibitory synapses. Eur. J. Cell Biol. 83, 449–456.

Vicidomini, G., Moneron, G., Han, K.Y., Westphal, V., Ta, H., Reuss, M., Engelhardt, J., Eggeling, C., and Hell, S.W. (2011). Sharper low-power STED nanoscopy by time gating. Nat. Methods 8, 571–573.

Wang, J., Tao, Y., Song, F., Sun, Y., Ott, J., and Saffen, D. (2015). Common Regulatory Variants of *CYFIP1* Contribute to Susceptibility for Autism Spectrum Disorder (ASD) and Classical Autism. Ann. Hum. Genet. 79, 329–340.

White, J.K., Gerdin, A.-K., Karp, N.A., Ryder, E., Buljan, M., Bussell, J.N., Salisbury, J., Clare, S., Ingham, N.J., Podrini, C., Houghton, R., Estabel, J., Bottomley, J.R., Melvin, D.G., Sunter, D., Adams, N.C., Tannahill, D., Logan, D.W., Macarthur, D.G., et al. (2013). Genome-wide generation and systematic phenotyping of knockout mice reveals new roles for many genes. Cell 154, 452–464.

Woo, J., Kwon, S.-K., Nam, J., Choi, S., Takahashi, H., Krueger, D., Park, J., Lee, Y., Bae, J.Y., Lee, D., Ko, J., Kim, H., Kim, M.-H., Bae, Y.C., Chang, S., Craig, A.M., and Kim, E. (2013). The adhesion protein IgSF9b is coupled to neuroligin 2 via S-SCAM to promote inhibitory synapse development. J. Cell Biol. 201, 929–944.

Xu, C., Fu, X., Zhu, S., and Liu, J.-J. (2016). Retrolinkin recruits the WAVE1 protein complex to facilitate BDNF-induced TrkB endocytosis and dendrite outgrowth. Mol. Biol. Cell 27, 3342–3356.

Yamasaki, T., Hoyos-Ramirez, E., Martenson, J.S., Morimoto-Tomita, M., and Tomita, S. (2017). GARLH family proteins stabilize GABAA receptors at synapses. Neuron 93, 1138–1152.e6.

Yizhar, O., Fenno, L.E., Prigge, M., Schneider, F., Davidson, T.J., O’Shea, D.J., Sohal, V.S., Goshen, I., Finkelstein, J., Paz, J.T., Stehfest, K., Fudim, R., Ramakrishnan, C., Huguenard, J.R., Hegemann, P., and Deisseroth, K. (2011). Neocortical excitation/inhibition balance in information processing and social dysfunction. Nature 477, 171–178.

Yoon, K.-J., Nguyen, H.N., Ursini, G., Zhang, F., Kim, N.-S., Wen, Z., Makri, G., Nauen, D., Shin, J.H., Park, Y., Chung, R., Pekle, E., Zhang, C., Towe, M., Hussaini, S.M.Q., Lee, Y., Rujescu, D., St Clair, D., Kleinman, J.E., et al. (2014). Modeling a Genetic Risk for Schizophrenia in iPSCs and Mice Reveals Neural Stem Cell Deficits Associated with Adherens Junctions and Polarity. Cell Stem Cell 15, 79–91.

Zhang, B., Seigneur, E., Wei, P., Gokce, O., Morgan, J., and Südhof, T.C. (2016). Developmental plasticity shapes synaptic phenotypes of autism-associated neuroligin-3 mutations in the calyx of Held. Mol. Psychiatry 22, 1483–1491.

van der Zwaag, B., Staal, W.G., Hochstenbach, R., Poot, M., Spierenburg, H.A., de Jonge, M. V, Verbeek, N.E., van ‘t Slot, R., van Es, M.A., Staal, F.J., Freitag, C.M., Buizer-Voskamp, J.E., Nelen, M.R., van den Berg, L.H., van Amstel, H.K.P., van Engeland, H., and Burbach, J.P.H. (2010). A co-segregating microduplication of chromosome 15q11.2 pinpoints two risk genes for autism spectrum disorder. Am. J. Med. Genet. B. Neuropsychiatr. Genet. 153B, 960–966.

## Supplemental References

López-Doménech, G., Higgs, N.F., Vaccaro, V., Roš, H., Arancibia-Cárcamo, I.L., MacAskill, A.F., and Kittler, J.T. (2016). Loss of Dendritic Complexity Precedes Neurodegeneration in a Mouse Model with Disrupted Mitochondrial Distribution in Mature Dendrites. Cell Rep. 17, 317–327.

Vaccaro, V., Devine, M.J., Higgs, N.F., and Kittler, J.T. (2017). Miro1-dependent mitochondrial positioning drives the rescaling of presynaptic Ca2+ signals during homeostatic plasticity. EMBO Rep. 18, 231–240.

